# Missing taxane C13α-*O*-deacetylases reveal a cryptic acetylation–deacetylation module in paclitaxel biosynthesis

**DOI:** 10.64898/2026.04.28.721278

**Authors:** Changkang Li, Xincheng Sun, Ridao Chen, Kebo Xie, Dawei Chen, Jimei Liu, Jungui Dai

**Affiliations:** State Key Laboratory of Bioactive Substance and Function of Natural Medicines; NHC Key Laboratory of Biosynthesis of Natural Products; and Beijing Key Laboratory of Key Technologies for Preclinical Research and Development of Innovative Drugs in Pharmacokinetics and Pharmacodynamics; Institute of Materia Medica, Chinese Academy of Medical Sciences and Peking Union Medical College, Beijing 100050, China

## Abstract

The prevalence of naturally occurring C13-acetoxy taxanes, together with the presence of a native C13-acetyltransferase in yew trees, suggests that the natural biosynthetic pathway for paclitaxel may involve a cryptic C13α-*O*-deacetylation step. However, whether a putative taxane C13α-*O*-deacetylase (T13dA) acts in the pathway of paclitaxel biosynthesis remains elusive. Herein, we functionally characterized two novel taxane C13α-*O*-deacetylases (T13dA1 and T13dA2) from *Taxus × media* cell cultures, providing experimental evidence for the molecular and biochemical plausibility of C13α-*O*-deacetylation in paclitaxel biosynthesis in *Taxus* species. Furthermore, T7dA1, a novel taxane C7β-*O*-deacetylase with higher activity than the reported T7dA was discovered and characterized here. Notably, we reconstituted a new 19-gene pathway enabled by the integration of a C13α-*O*-acetylation-deacetylation module for the *de novo* biosynthesis of baccatin III in *Nicotiana benthamiana* leaves, reaching a yield of 71**□** μg**□** g^−1^ DW, much higher than that obtained from the 17**□** gene pathway lacking this module. These findings not only demonstrate the functional importance of the C13α-*O*-acetylation-deacetylation module as a branch within the paclitaxel biosynthetic network, but also expand the multi-branched taxane metabolic network in *Taxus* and provide novel enzymatic tools and engineering strategies to facilitate heterologous pathway reconstitution and efficient paclitaxel production.

## Introduction

Paclitaxel (trade name Taxol), a clinically potent anticancer drug derived from *Taxus* species, features a unique 6/8/6-membered carbocyclic diterpene scaffold with extensive oxygenation and complex functionalities^1^. However, plant-derived paclitaxel suffers from a short supply due to its low abundance in *Taxus* together with inefficient total chemical synthesis, leading to persistent challenges in sustainable supply^2^. The current paclitaxel supply depends on costly semi-synthesis, in which paclitaxel precursor, baccatin III, is isolated from raw tree biomass or plant cell cultures of *Taxus* and subsequently chemically converted into paclitaxel^3,4^. However, the semi-synthesis strategy relies heavily on natural resources and artificial cultivation of *Taxus* plants^5^. With the advancement of synthetic biology, the biosynthetic strategy for green and sustainable paclitaxel production has become increasingly appealing^6,7^. For the foreseeable future, the efficient and affordable supply of paclitaxel and its precursor baccatin III will rely upon a synthetic biology approach, which necessitates a better understanding of the biosynthesis of paclitaxel and related taxoids in nature, and the discovery of the responsible enzymes and components^8,9^.

Generally, the paclitaxel biosynthetic pathway has been characterized biochemically and can be divided into three stages (Supplementary Fig. S1a)^10^: (1) the taxadiene formation begins with geranylgeranyl pyrophosphate (GGPP)^11^, a key common precursor of diterpenoids, (2) conversion of taxadiene to baccatin III (**1**) and 10-deacetylbaccatin III (10-DAB, **2**)^3,12^, and (3) attachment of C13-side chain to baccatin III or 10-DAB^3^, which forms paclitaxel (**3**). By 2025, multiple groups have discovered 23 enzymes involved in paclitaxel biosynthesis^7,13^, including the scaffold-forming enzyme, taxadiene synthase (TXS)^11^, a variety of tailoring oxidases (T5αH^14^, T13αH^15^, T10βH^16^, T9αHs (T9αH-725A^17^ and T9αH-750C^12^), T7βH^18^, T2αH^19^, T1βH^12,20^, TOT^10,21,22^, T9Ox^17^), four acyltransferases (TAT^23^, DBAT^24^, T7AT^22^, TBT^25^), two taxane deacetylases (T7dA, and T9dA)^12^, as well as a facilitator of taxane oxidation (FoTO1)^12^, which collectively generate a key paclitaxel precursor baccatin III. The C13-side chain is subsequently attached to baccatin III by five additional enzymes (PAL, TBPCCL, BAPT, T2**′**OGD, T3**′**NBT) to yield paclitaxel^3^. The biosynthetic C13-side chain formation and its combination with baccatin III to produce paclitaxel have been well studied, and baccatin III (**1**) is considered as the target product for heterologous production because of the chemically easy accessibility to the industrial C13-side chain and further coupling with baccatin III. Up to now, three groups have achieved the *de novo* biosynthesis of baccatin III in *N. benthamiana* leaves without optimization. Zhang^17^ and Jiang^10^ reconstituted a 13-gene set of baccatin III pathway (Supplementary Fig. S1b, route i) and a 9-gene set baccatin III pathway in tobacco system (Supplementary Fig. S1b, route ii)^10^, respectively. However, these two published gene sets lacked several biosynthetic enzymes, resulting in a low yield of baccatin III; its production may be attributable to the multifunctionality of certain *Taxus* enzymes or to that of endogenous *N. benthamiana* enzymes. Subsequently, Sattely and coworkers discovered the missing key enzyme T1βH, and employed the two new functionally characterized downstream deacetylases T7dA and T9dA to build a complete 17-gene pathway (Fig. 1) of baccatin III in tobacco^12^. Besides, the T9αH used here is a CYP750C enzyme instead of a CYP725A enzyme. More importantly, C7- and C9-deacetylation steps were first demonstrated to be involved in the biosynthesis of paclitaxel, and considered as protection for the next oxidation(s).

**Fig. 1.**
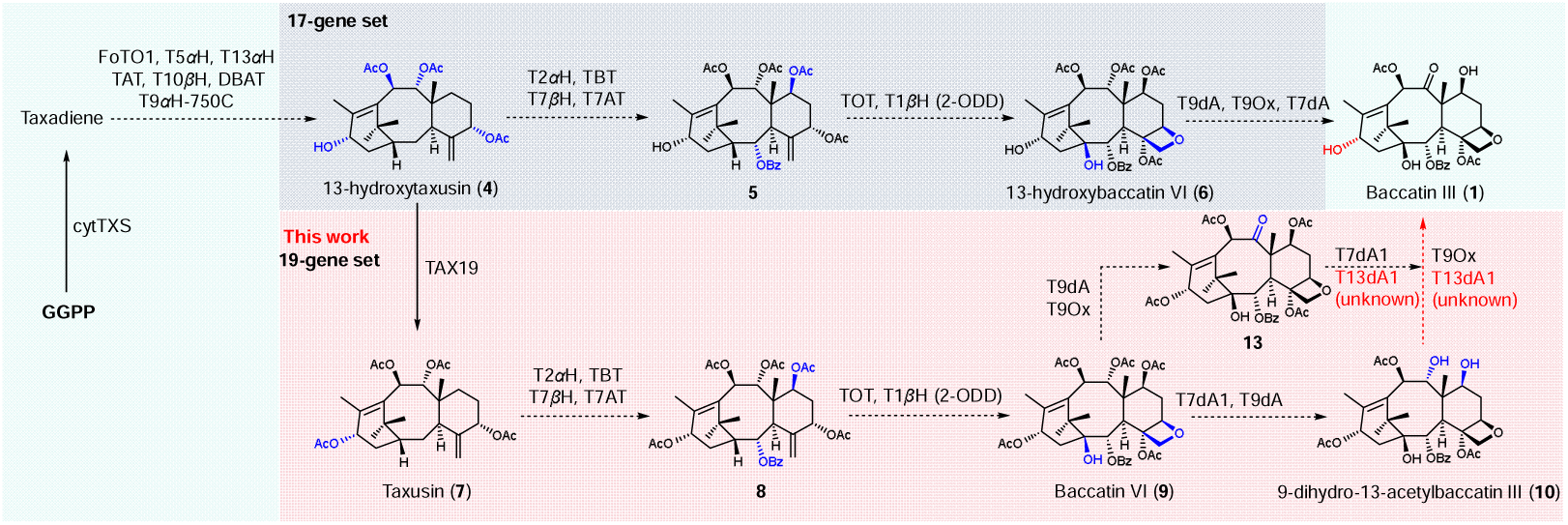
The previously published 17-gene pathway and heterologous reconstitution of baccatin III in *Nicotiana benthamiana* leaves without the C13-acetylation-deacetylation module, and the 19-gene pathway that includes this module, which is the focus of the present study. This pathway provides the first direct biochemical evidence that deacetylation steps are functionally required component of paclitaxel biosynthetic pathway. A defining characteristic of this pathway is the consistent presence of a hydroxyl group at C13 position. FoTO1: facilitator of taxane oxidation, a nuclear transport factor 2 (NTF2)-like protein; TXS: taxadiene synthase; cytTXS: TXS with an N-terminal truncation removing the native chloroplast localization sequences; T5αH: taxane C5α-hydroxylase; T13αH: taxane C13α-hydroxylase; T10βH: taxane C10β-hydroxylase; T9αH-750C: taxane C9α-hydroxylase belonging to CYP750C enzyme (cytochrome P450, family 750, subfamily C); T7βH: taxane C7β-hydroxylase; T2αH: taxane C2α-hydroxylase; T9Ox: taxane C9α-*O*-oxidase (oxidation of the C9-hydroxy to the corresponding carbonyl); TAT: taxane C5α-*O*-acetyltransferase; TBT: taxane C2α-*O*-benzoyltransferase; DBAT: taxane C10β-*O*-acetyltransferase; TOT: taxane oxetanase; T1βH (2-ODD): taxane C1β-hydroxylase belonging to 2-oxoglutarate dependent dioxygenase; TAX19: taxane C13α-*O*-acetyltransferase. T7dA: taxane C7β-*O*-deacetylase; T9dA: taxane C9α-*O*-deacetylase; T13dA: taxane C13α-*O*-deacetylase (not yet functionally characterized).

The above discovery of two taxane deacetylases (T7dA and T9dA) indicated the occurrence of cryptic acetylation-deacetylation in the biosynthesis of paclitaxel. To our knowledge, installation of C13α-hydroxyl group occurs at an early stage (the first two hydroxylation steps) of paclitaxel biosynthesis^15^, and the presence of TAX19^23^, a previously characterized taxane C13α-*O*-acetyltransferase from *Taxus*, suggests that a cryptic C13-acetylation-deacetylation process is invovled in the biosynthesis of paclitaxel. In addition, structural characterization of several hundred defined taxoids bearing oxy-functional groups at various positions isolated from *Taxus* species reveals that most natural taxanes possess an acetoxy group at the C13 position^26,27^, further supporting the above deduction. In our recent work^20^, we reconstituted the *de novo* biosynthetic pathway of baccatin VI (**9**) which has three additional acetyl groups (C7β-*O*-, C9α-*O*-and C13α-*O*-) in tobacco. However, whether a dedicated C13α-O-deacetylase (T13dA) and a cryptic acetylation-deacetylation in the transformation of baccatin VI (**9**) to baccatin III (**1**) exist has remained unclear.

In this work, we functionally characterized two missing taxane C13α-*O*-deacetylases (T13dA1 and T13dA2) from the cell cultures of *T. × media*, alongside a novel taxane C7β**□** *O***□** deacetylase (T7dA1) variant that exhibits markedly higher enzymatic activity than the previously characterized T7dA^12^. These three enzymes act as the previously uncharacterized deacetylases within a distinct sub**□** branch of the paclitaxel biosynthetic network. Notably, the discovery of T13dA enzymes reveals a cryptic acetylation–deacetylation module that has not been recognized in the canonical pathway. Based on these findings, we reconstituted a 19**□** gene *de novo* baccatin III pathway with the C13-acetylation–deacetylation module in tobacco, achieving a maximum yield of 71**□** μg**□** g^-1^ DW, substantially higher than that of the 17**□** gene pathway without this module. These results demonstrate that this module constitutes a functionally important sub-branch of the paclitaxel biosynthetic metabolic network and provide new insights into the multi-branched taxane metabolic pathways in *Taxus* species.

## Results and discussion

### The discovery of missing taxane deacetylases involved in paclitaxel biosynthesis

Our latest reconstituted the pathway of intermediate baccatin VI (**9**) from GGPP in tobacco has three additional acetylations at C7, C9 and C13 positions that are not found in baccatin III^20^. Given the known taxane deacetylases T7dA and T9dA can stepwise remove the C7-*O*- and C9-*O*-acetyl groups from **9** to generate 9-dihydro-13-acetylbaccatin III (**10**, Fig 1), we hypothesized that **10** undergoes further conversion to baccatin III (**1**) through the combined action of a putative taxane C13α-*O*-deacetylase (T13dA) and the known oxidase T9Ox (Fig. 1). Therefore, both compounds **9** and **10** could be used as candidate substrates for the discovery and functional identification of the missing T13dA.

Carboxylesterase, a key member of the α/β-hydrolase superfamily^28,29^, plays a pivotal role in the deacetylation during the biosynthesis of plant-derived natural products, such as noscapine^30^ and azadirone^31^. Therefore, we focused our search on carboxylesterase to mine the missing T13dAs, using RNA-sequencing data from paclitaxel-producing *T. × media* cell cultures for gene discovery^20,21^. As a result, fifty-three full-length candidate genes were obtained by screening the transcriptomic dataset with the keyword “carboxylesterase”. We first opted to use *N. benthamiana* transient expression system for rapid screening of active enzymes. All fifty-three candidate genes were then divided into five groups (A–E) *via* phylogenetic analysis (Fig. 2a) and simultaneously expressed all the genes in each group in *N. benthamiana* leaves using *Agrobacterium*-mediated transformation. After four days of cultivation, substrates **9** and **10** were individually fed into transgenic tobacco leaves, which were collected after an additional 24 h of incubation. The leaves were extracted with methanol after lyophilization for 48 h and analysed by liquid chromatography–mass spectrometry (LC–MS).

**Fig. 2.**
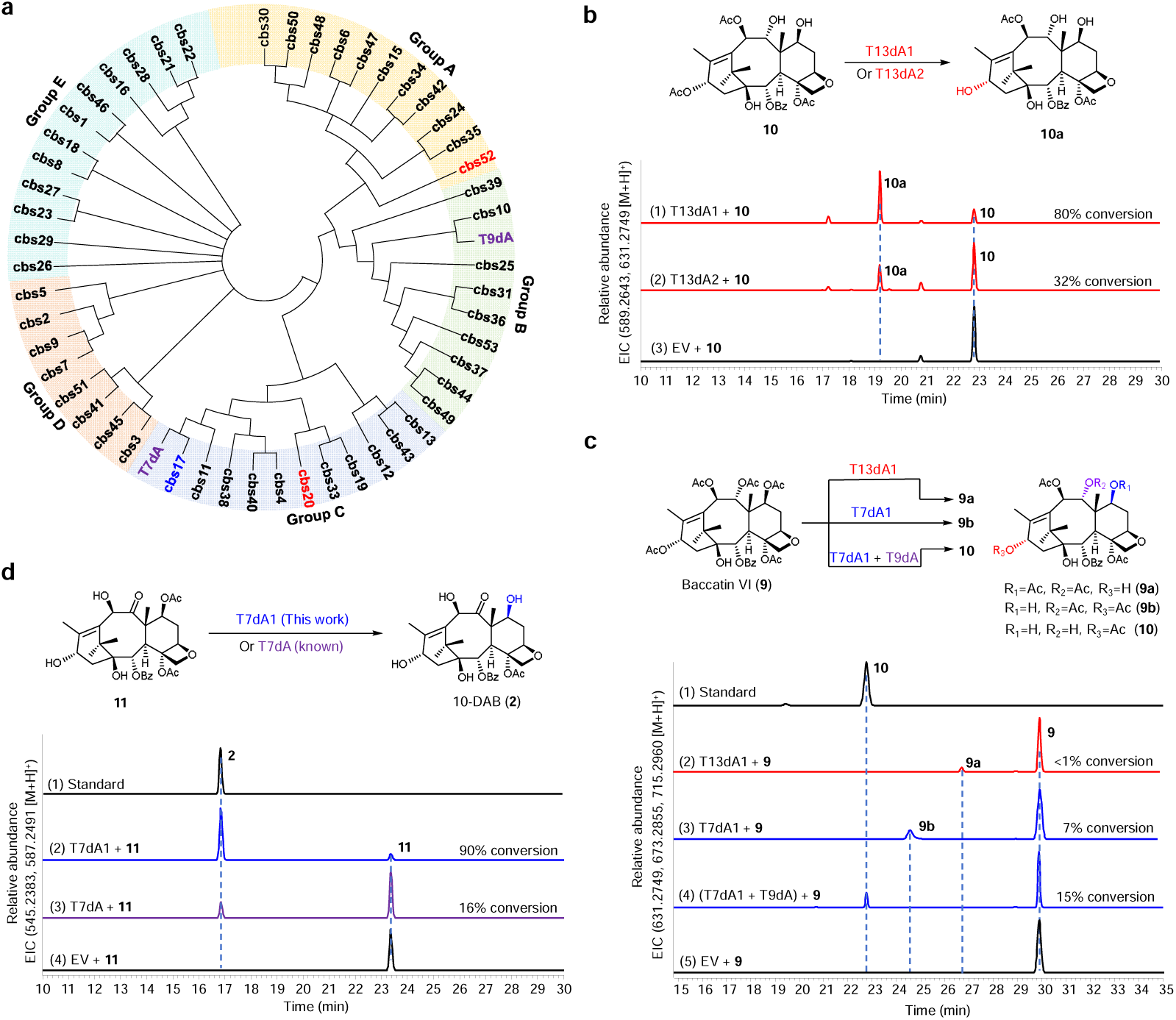
Discovery and functional characterization of three taxane deacetylases, T13dA1, T13dA2 and T7dA1. **a**, Phylogenetic analysis of carboxylesterase in *Taxus × media* cell cultures. The phylogenetic trees were constructed using the neighbor-joining method in Mega with 1000 bootstrap replicates. **b**, Functional characterization of T13dA1 and T13dA2. Either T13dA1 or T13dA2 is capable of independently and efficiently catalysing the conversion of **10** to **10a** in *Nicotiana benthamiana* leaves. Extracted ion chromatograms (EICs, [M + H]^+^) of extracts from *N. benthamiana* leaves with transient expression of (1) T13dA1 + **10**, (2) T13dA2 + **10**, and (3) empty vector (EV) + **10**. **c**, Functional characterization of T13dA1 and T7dA1 towards substrate **9** in *N. benthamiana* leaves. Products **9a** and **9b** were generated by T13dA1- and T7dA1-catalyzed deacetylation of substrate **9**, respectively. Product **10** corresponds to the sequential deacetylation of substrate 9 by the enzyme combination T7dA1 + T9dA. Shown are EICs ([M + H]^+^) of products **9a**, **9b** (*m/z* 673.2855), **10** (*m/z* 631.2749) and substrate **9** (*m/z* 715.2960). **d**, Comparison of T7dA1 and T7dA activities toward substrate **11** in *N. benthamiana* leaves. EICs ([M + H]^+^) of extracts from *N. benthamiana* leaves with transient expression of (1) standard of **12**; (2) T7dA1 + **11**; (3) T7dA + **11**; and (4) empty vector (EV) + **1**.

As a result, the HPLC peak area of substrate **10** in the groups A and C was considerably decreased, respectively, yielding two dominant peaks with the identical retention time, whose mass features correspond to a mono-deacetylated product (**10a**, Supplementary Fig. S2a). MS/MS fragmentation patterns indicated that the two chromatographic peaks corresponded to the same compound (Supplementary Fig. S2b), suggesting that both group A and group C contain enzymes capable of catalyzing the same deacetylation reaction of **10**. To determine which genes in groups A and C could catalyse the substrate **10**, we individually performed substrate-feeding experiments by expressing each gene in the two groups in tobacco leaves. We observed that two of these candidates (cbs52 from group A and cbs20 from group C) led to the production of **10a** with high conversion (Fig. 2b). Interestingly, cbs52 and cbs20 share only 7% sequence identity at the protein level and showed the same reactivity towards **10** in tobacco system. Moreover, cbs52 results in significantly higher conversion rate of the deacetylated product **10a** (ca. 80% conversion) than cbs20 (ca. 32% conversion), thus, cbs52 was used in all subsequent experiments.

Considering that carboxylesterase can be expressed in a soluble form, cbs52 was heterologously expressed as N-terminal His_6_-tagged proteins in *Escherichia coli* Transetta DE3. Purification of protein cbs52 was accomplished by His-tag affinity chromatography and analysis by SDS-PAGE (Supplementary Fig. S3). To characterize the structure of product **10a**, we further performed *in vitro* enzymatic reactions at large scale and isolated the resulting deacetylated product **10a**. Through 1D/2D NMR and HR-ESI-MS spectroscopic analysis, product **10a** was structurally characterized as 9-dihydrobaccatin III, a C13-deacetylated product (Fig. 2b). These results indicate that cbs52 is a deacetylase responsible for taxane C13α-*O*-deacetylation, referred to as T13dA1 (GenBank accession no. PX848827), and isoenzyme cbs20 was named T13dA2 (GenBank accession no. PX848828).

However, in addition to T13dA1 (from group A) being capable of converting substrate **9** to **9a** with low levels (Fig 2c), we observed that cbs17 (from group C) catalyzes the *mono*-deacetylation of substrate **9** to yield product **9b** (Fig. 2c). Its specific catalytic function was unequivocally confirmed by feeding chemically synthesized taxoid 7-acetoxy-10-deacetylbaccatin III (**11**) to *N. benthamiana* leaves transiently expressing cbs17; the enzyme removed the C7-acetyl group from **11** to produce a product matching the 10-deacetylbaccatin III (10-DAB, **2**) standard (Fig. 2d), as supported by MS/MS analysis (Supplementary Fig. S4). Therefore, cbs17 is classified as a novel taxane C7β-*O*-deacetylase and designated T7dA1 (GenBank accession no. PX848829). Phylogenetic analysis showed that T7dA1 is closely related to the known T7dA (Fig. 2a). Moreover, tobacco feeding assays verified that the enzyme pair (T7dA1 + T9dA) catalyzes deacetylation at the C7 and C9 sites of compound **9**, generating a product identical to the authentic standard of compound **10** (Fig. 2c), which was validated *via* MS/MS spectral comparison with the reference standard (Supplementary Fig. S5). This result further substantiates the C7 deacetylation function of T7dA1. Notably, T7dA1 was shown to efficiently catalyse C7-deacetylation of **11**, exhibiting substantially higher catalytic activity than the previously reported variant T7dA (Fig. 2d). Therefore, T7dA1 was used as the taxane C7β-*O*-deacetylase in all subsequent experiments.

Next, T7dA1 was also heterologously expressed in *E. coli*, and the corresponding proteins were purified (Supplementary Fig. S3). The biochemical properties of the three taxane deacetylases (T13dA1 and T7dA1) were examined. Different pH values and temperatures were assayed to optimize the deacetylase reaction conditions. The optimal pH for T13dA1 activity was pH 8.0 (50 mM Tris-HCl buffer Supplementary Fig. S6). The optimal pH for T7dA1 was pH 7.0 (50 mM NaH_2_PO_4_-Na_2_HPO_4_ buffer, Supplementary Fig. S6). Under different tested temperatures, T13dA1 and T7dA1 exhibited the highest catalytic activity at 30°C and 35°C, respectively (Supplementary Fig. S6).

### Integration of a cryptic C13α-*O*-acetylation–deacetylation module into the paclitaxel biosynthetic pathway

Having identified T13dA1 as the enzyme catalysing the deacetylation steps of paclitaxel pathway, we next attempted to delineate a novel paclitaxel biosynthesis pathway featuring C13-acetylation/deacetylation steps. In the previously reported 17-gene baccatin III pathway without C13-acetylation–deacetylation involvement^12^, the C13 position is consistently substituted with a hydroxyl group, among which C13-hydroxylated products 13-hydroxytaxusin (**4**) and 13-hydroxybaccatin VI (**6**) serve as key intermediates (Fig. 1). By contrast, the introduction of C13-acetylation and deacetylation steps into the above 17-gene baccatin III pathway would likely result in the accumulation of C13-acetylated products taxusin (**7**) and baccatin VI (**9**) as the predominant biosynthetic intermediates (Fig. 1). To verify this hypothesis, we fed taxusin (**7**) and baccatin VI (**9**) into *N. benthamiana* leaves co-expressing 10-gene combination (T2αH + T7βH + TBT + T7AT + TOT + T1βH + T7dA1 + T9dA + T13dA1 + T9Ox, combination 1) and 4-gene combination (T7dA1 + T9dA + T13dA1 + T9Ox, combination 3), respectively. As expected, both **7** and **9** can be transformed into baccatin III (**1**) when the C13α-*O*-deacetylase (T13dA1) was introduced into baccatin III pathway (Fig. 3a, Supplementary Fig. S7). As a control, omission of T13dA1 (Fig. 3a, combinations 2 and 4) abolished baccatin**□** III production while 3α**□** acetoxybaccatin**□** III (**12**) could be detectable. This result demonstrates that the conversion to baccatin**□** III is strictly dependent on T13dA1 and excludes interference from endogenous background deacetylase activity in *N. benthamiana*. In addition, baccatin VI (**9**) was detected in combination 1 pathway (Fig 3a), confirming its role as an intermediate in the conversion of taxusin (**7**) to baccatin III (**1**). Notably, we observed that the production of baccatin III (**1**) starting from taxusin (**7**) was markedly greater than that from baccatin VI (**9**). This difference likely indicates the availability of multiple alternative routes from taxusin (**7**) to baccatin III (**1**), further suggesting that paclitaxel biosynthesis is better described as a metabolic network than a simple linear sequence. However, the pathway we proposed merely represents one branch supported by our current experimental evidence, and more optimal catalytic routes might exist for further exploration. Additionally, such efficiency discrepancy suggests that upstream enzymes might exert scaffolding functions beyond their core catalytic activity to facilitate downstream enzymatic conversions.

**Fig. 3.**
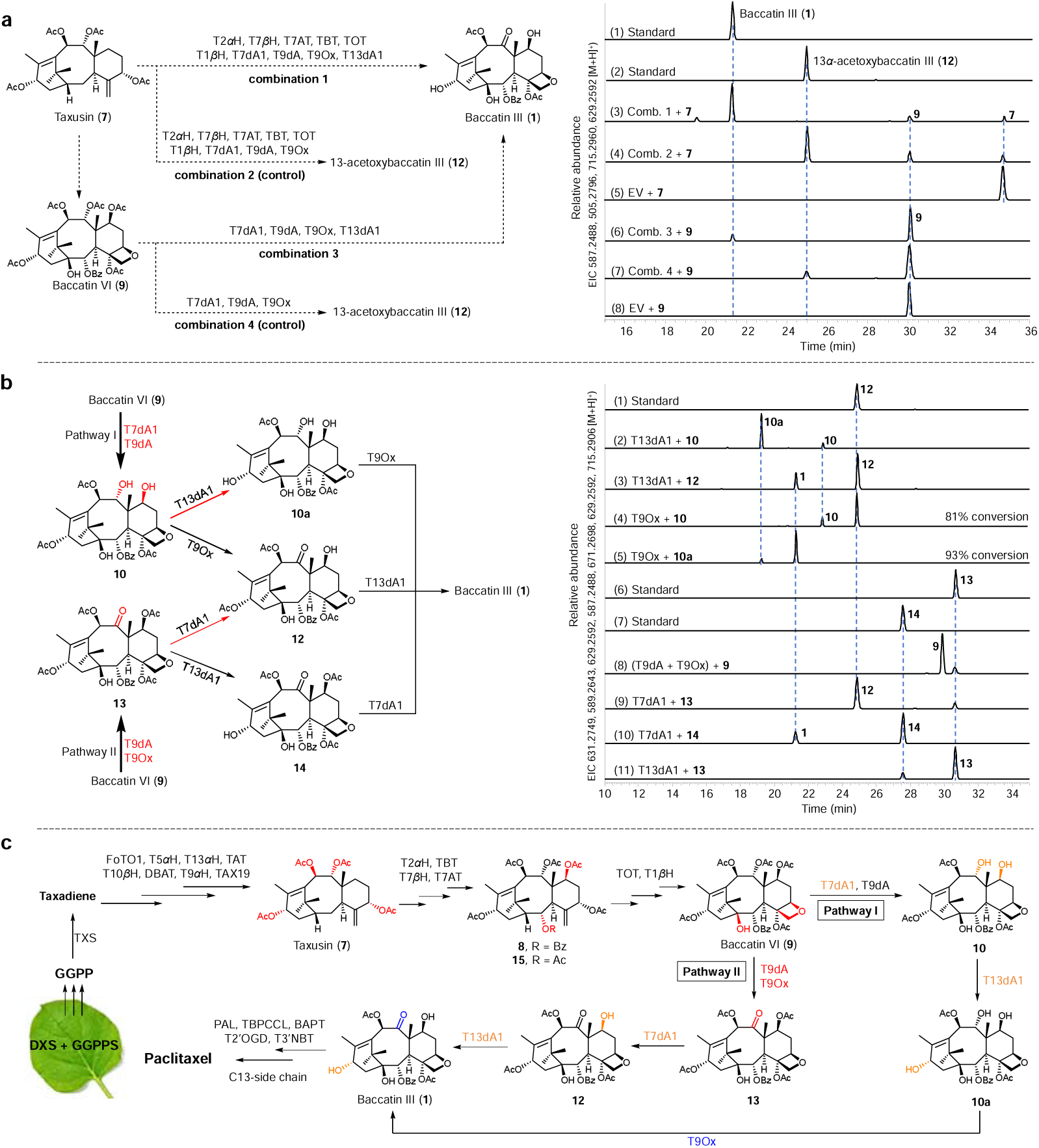
Missing taxane deacetylases in a new biosynthetic pathway for baccatin III. **a**, Proposed biosynthetic pathway from taxusin (**7**) and baccatin VI (**9**) to baccatin III (**1**). LC–MS chromatograms show the formation of **1** from **7** and **9** through the transient pathway expression (combinations 1 and 3) in *N. benthamiana* leaves, compared to an authentic standard of **1**. Baccatin III (**1**) was detected in combinations 1 and 3. By contrast, no baccatin III (**1**) was observed in conmbinations 2 and 4 without T13dA1, whereas 13α-acetoxybaccatin III (**12**) accumulated in these two control groups instead. Baccatin VI (**9**) was detected in combination 1 pathway. **b**, The specific catalytic orders and substrate promiscuities of T13dA1, T7dA1 and T9Ox. Red arrows represent the predominant pathway. EICs of extracts from *N. benthamiana* leaves with transient expression of (2) T13dA1 + **10**, (3) T13dA1 + **12**, (4) T9Ox + **10**, (5) T9Ox + **10a**, (8) (T9dA+T9Ox) + **9**, (9) T7dA1 + **13**, (10) T7dA1 + **14**, (11) T13dA1 + **13** compared to that of **12**−**14** standards. **c**, Re-delineated a novel paclitaxel biosynthetic pathway involving a cryptic C13-acetylation-deacetylation module. The enzymes identified in this study are highlighted in orange.

Several studies have demonstrated that the intermediate 2-benzoyloxy-5,7,9,10,13-pentaacetyloxytax-4(20),11-diene (**8**, Fig. 1) can serve as a direct substrate for TOT enzyme^22^, enabling oxetane ring formation, followed by T1βH-mediated C1β-hydroxylation, to afford baccatin VI (**9**)^12,20^. Subsequently, starting from baccatin VI (**9**), we proposed two plausible biosynthetic pathways (pathways I and II, Fig. 3b) and experimentally confirmed their feasibility through enzymatic reactions of key intermediates. As previously demonstrated (Fig. 2c), the deacetylation reactivity of baccatin VI at the C13 position is significantly lower (<1% conversion) than that at the C7 and C9 positions. Therefore, in pathway I, deacetylation at the C7 and C9 of baccatin VI (**9**) takes priority. We have experimentally demonstrated that the combination of (T7dA1 + T9dA) confers considerable catalytic efficiency for the deacetylation of baccatin VI (**9**), acting sequentially to mediate deacetylation at the C7 and C9 positions and yielding compound **10** (Fig. 2c). Finally, intermediate **10** is converted to baccatin III (**1**) *via* deacetylation at C13 and ketone formation at C9, catalyzed by T13dA1 and T9Ox. To investigate the sequence of the final two biosynthetic events, we synthesized the intermediate **12** bearing a C9 ketone functionality. *In vitro* steady-state kinetics indicated that T13dA1 had a much higher catalytic efficiency with **10** than with **12** (*k_cat_*/*K*_m_ **□**=**□** 114.2 ± 1.6**□**min^−1^mM^−1^ for **10** compared with 2.3 ± 0.1**□**min^−1^mM^−1^ for **12**, Supplementary Fig. S8ab), implying that T13dA1 prefers the substrate **10** for C13-deacetylation (Fig. 3b). Similarly, compared with substrate **10**, T9Ox prefers to accept **10a** as a substrate for oxidation of C9-hydroxy (Fig. 3b). The above data showed that the C13-deacetylation occurs prior to the oxidation of C9-OH. In fact, both T13dA1 and T9Ox exhibited appreciable catalytic activity toward these substrates, collectively contributing to the metabolic network of late paclitaxel pathway (Fig. 3b).

For pathway II, the C9-ketone formation was supposed to occur preferentially. To test this hypothesis, we infiltrated *N. benthamiana* leaves co-expressing T9dA and T9Ox with baccatin VI (**9**). A product matching the authentic standard of 7,13-diacetoxybaccatin III (**13**) was detected (Fig. 3b), which verifies that biosynthetic pathway II is functionally operative. Next, the intermediate **13** can be transformed into baccatin III (**1**) through deacetylation at the C7 and C13 by the action of T7dA1 and T13dA1, respectively. To investigate the sequence of these two deacetylation events, we synthesized intermediate **14** bearing a C13-hydroxy group. *In vitro* steady-state kinetics indicated that T7dA1 had a significant higher catalytic efficiency with **13** than with **14** (*k_cat_*/*K*_m_ **□** =**□** 36.8 ± 0.4**□**min^−1^mM^−1^ for **13** compared with 4.8 ± 0.2**□**min^−1^mM^−1^ for **14**, Supplementary Fig. S8cd), implying that T7dA1 prefers the substrate **13** for C7-deacetylation (Fig. 3b). In contrast, T13dA1 exhibits comparable catalytic activity toward substrates **13** and **12**, with an overall activity marginally lower than that of T7dA1 (Fig. 3b). These findings indicate that introduction of the C9-ketone prior to deacetylation results in preferential deacetylation at the C7 position relative to the C13 position (pathway II).

Accordingly, by integrating the cryptic C13α**□***O***□**acetylation–deacetylation module into the metabolic framework of paclitaxel biosynthesis, we propose a novel biosynthetic route for paclitaxel. (Fig. 3c): The diterpenoid precursor GGPP is first cyclized by taxadiene synthetase (TXS), generating a unique diterpene skeleton, taxadiene, followed by hydroxylation and acetylation at C5, C10, C13, and C9 of taxadiene to yield taxusin (**7**), which is generally regarded as the early stage of paclitaxel biosynthesis. Subsequently, hydroxylation and acylation occur at C2 and C7 of taxusin (**7**), yielding 2-benzoyloxy-5,7,9,10,13-pentaacetoxytaxadiene (**8**). Interestingly, several studies have demonstrated that the taxane C2-*O*-benzoyltransferase (TBT) can also mediate the generation of acetylated product (**15**) in tobacco, which may be diverted into a non-paclitaxel pathway. This bifunctionality might explain the prevalence of C2α-acetoxy taxanes in nature^12,20^. Next, intermediate **8** undergoes the oxetane ring formation and C1β-hydroxylation to yield baccatin VI (**9**). Starting from baccatin VI (**9**), the biosynthetic route diverges into two distinct pathways: pathway I, characterized by preferential C7- and C9-deacetylation, and pathway II, defined by initial C9 ketone formation. In pathway I, T7dA1 and T9dA act cooperatively the deacetylation of baccatin VI (**9**) at C7 and C9 positions to yield intermediate **10**. Intermediate **10** then undergoes sequential C13-deacetylation and C9-ketone formation to afford baccatin III (**1**). In pathway II, the C9 ketone is preferentially formed to yield intermediate **13** under the concerted catalysis of T9dA and T9Ox; subsequently, intermediate **13** undergoes sequential deacetylation at C7 and C13 positions to afford baccatin III. Finally, the C13-side chain is attached to baccatin III to generate paclitaxel.

### Heterologous production and quantitative analysis of baccatin III in *N. benthamiana*

In our previous work^20^, the biosynthetic pathway from GGPP to baccatin VI was reconstituted in *N. benthamiana* system, but failed to produce the expected baccatin III (**1**), possibly attributable to the absence of key functional taxane deacetylases, thereby blocking the conversion of baccatin VI to baccatin III. In this study, we functionally identified three novel deacetylases (T13dA1 and T13dA2, and T7dA1). Therefore, we attempted to integrate these enzymes into the aforementioned baccatin VI pathway to enable *de novo* production of baccatin III in *N. benthamiana*. To the best of our knowledge, plants synthesize terpenoids *via* two distinctly compartmentalized endogenous pathways: (1) the cytosol-localized mevalonate (MEV, Fig. 4a) pathway, and (2) the chloroplast-localized 1-deoxy-D-xylulose-5-phosphate (DXP, Fig. 4a) pathway^32−34^. The isopentenyl diphosphate (IPP) derived from the plastid DXP pathway is employed in the biosynthesis of diterpenes^33−35^. Previous reports have shown that 1-deoxy-D-xylulose-5-phosphate synthase (DXS) is a rate-limiting enzyme for plastidic isoprenoid biosynthesis in plants^36^. Overexpression of DXS and geranylgeranyl diphosphate synthase (GGPPS) could significantly increase the accumulation of geranylgeranyl diphosphate (GGPP), thereby facilitating the biosynthesis of diterpenoids in plants^37−40^. Based on this strategy, we first attempted to supply GGPP endogenously by co**□** expressing DXS and GGPPS in *N. benthamiana* as one of our approaches.

**Fig. 4.**
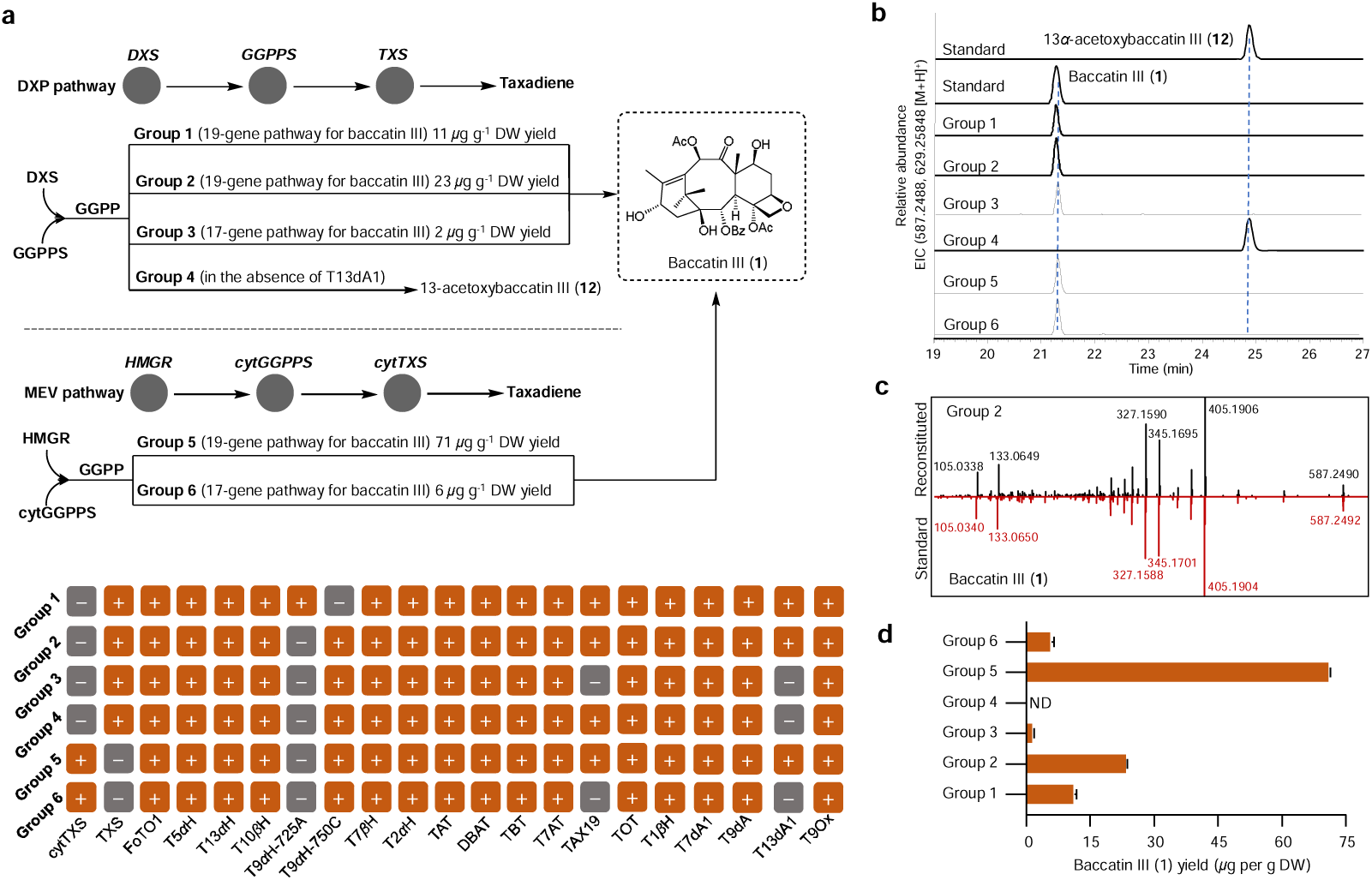
*De novo* biosynthesis of baccatin III in *N. benthamiana* and its quantitative analysis. **a**, Multiple enzyme combinations (groups 1−6) were employed for baccatin III pathway reconstruction. TAT is a bi-functional acetyltransferase that introduces the acetyl groups on both the C5α and C9ß hydroxyls^12^. TAX19: taxane C13α-*O*-acetyltransferase. GGPP was supplied via two distinct precursor pathways: Groups 1–4 employed the plastidic DXP route by co□expressing the rate□ limiting enzyme DXS and GGPPS, whereas Groups 5 and 6 used the cytosolic MEV route by co□expressing HMGR and cytGGPPS. HMGR: The 3□hydroxy□3□methylglutaryl□ CoA reductase (AY488113) employed in this study originates from *Arabidopsis thaliana*^32,41^. **b**, EICs for baccatin III (**1**) in tobacco leaves expressing different gene combinations (shown as groups 1−6). **c**, MS/MS spectrum of baccatin III produced in tobacco (black, group 2) and its standard (red). **d**, Bar chart depicting the baccatin III yield achieved with different gene combinations. Baccatin III (**1**) yields in μg per g dried weight (DW), quantified with standards (Methods). Data are mean□±□ s.d., *n*□=□3 biological replicates.

Upon integration of the C13 acetylation–deacetylation module, the new biosynthetic route toward baccatin III requires a total of 19 genes. Given that C9-hydroxylation is independently catalysed by two distinct T9αH enzymes originating from separate P450 families (CYP725A and CYP750C), we utilised T9αH**□**725A to validate baccatin III biosynthesis *via* transient expression in *N. benthamiana* (Fig. 4a, Group 1). This reconstitution system yielded baccatin III at approximately 11 μg g^-1^ dry weight (DW) (Fig. 4b), as confirmed by LC–MS comparison to an authentic standard (Fig. 4c, Supplementary Fig. S9). Next, to evaluate whether T9αH-750C maintains functional compatibility upon integration of the C13-acetylation–deacetylation module, we substituted T9αH-725A with T9αH-750C in group 1 (Fig. 4a, Group 2). As a result, the baccatin III yield increased to 23 μg g^-1^ DW, demonstrating that T9αH-750C exhibited superior performance relative to T9αH-725A after the introduction of C13-acetylation-deacetylation step into the paclitaxel biosynthetic pathway. In contrast, the absence of the C13-acetylation-deacetylation module in the pathway (Fig. 4a, Group 3) resulted in a marked decreasement in baccatin III production to 2 *µ*g g^-1^, demonstrating that this module plays a critical role in paclitaxel biosynthetic pathway. In the absence of T13dA1 (Fig. 4, Group 4), no baccatin**□** III production could be detected, whereas 13α**□** acetoxybaccatin**□** III (**12**) was detected (Fig. 4b, Supplementary Fig. S9), indicating that the C13-deacetylase is involved in the biosynthesis of baccatin**□**III.

However, Sattely and her colleagues demonstrated that redirecting the biosynthetic flux of diterpenoids from the low**□** flux plastidial DXP pathway to the high**□** flux cytosolic MEV pathway can substantially boost the yields of diterpene scaffolds^12,32^. Following this strategy, we removed the predicted N**□**terminal chloroplast transit peptides from both the target diterpene synthase (TXS) and geranylgeranyl pyrophosphate synthase (GGPPS), generating N**□** terminally truncated variants (designated cytTXS and cytGGPPS). These constructs were then co**□**expressed with the rate**□** limiting MEV pathway enzyme HMGR (3**□** hydroxy**□** 3**□** methylglutaryl**□**CoA reductase), thereby rerouting taxadiene production to the cytosol. To assess whether this rerouting strategy could improve the accumulation of baccatin III, we reconstituted the complete 19**□** gene baccatin III pathway in *N. benthamiana* leaves (Fig. 4a, Group 5, Supplementary Fig. S9). The final titer reached 71**□**μg**□** g^-1^ DW, representing a threefold increase compared with that obtained *via* the native DXP pathway. In contrast, when the C13 acetylation–deacetylation module was unincorporated, the 17**□** gene reconstituted pathway (Fig. 4a, Group 6, Supplementary Fig. S9) yielded 6**□** μg**□** g^-1^ DW. These results further indicate the functionally important branch of the C13 acetylation–deacetylation module in paclitaxel biosynthesis, and its engineering advantage for heterologous production.

## Conclusion

In this work, we identified three novel deacetylases in the paclitaxel biosynthetic pathway, including two previously missing taxane C13α-*O*-deacetylases (T13dA1 and T13dA2, Fig. 2b) responsible for taxane C13α-*O*-deacetylation, and a taxane C7β**□** *O***□**deacetylase (T7dA1, Fig. 2d) with markedly higher catalytic efficiency than the previously reported T7dA. The discovery of T13dAs reveals a cryptic acetylation-deacetylation module during paclitaxel biosynthesis. By integrating these newly identified deacetylases, we reconstituted a *de novo* 19-gene baccatin III biosynthetic pathway in *N. benthamiana* using a cytosolic MEV pathway engineering strategy^12,32^, achieving a maximum yield of 71 μg g^-1^ DW (Fig. 4, Group 5), over 10**□** fold higher than that with the 17**□** gene pathway without the C13 modification module (Fig. 4, Group 6). These results establish the C13 acetylation–deacetylation module as a functionally significant branch of paclitaxel biosynthesis.

Furthermore, the C13-acetylation–deacetylation module exerts a broader mechanistic influence on upstream pathway compatibility. Previous studies have demonstrated independent evolution of T9αH catalytic activities across two divergent P450 subfamilies (CYP725A and CYP750C), and their divergent substrate preferences imply that T9α**Η**-725A primarily supports the formation of C13α-acetoxyl taxanes, whereas T9αH-750C drives the biosynthesis of C13α-hydroxyl taxanes. In the conventional 17-gene baccatin III pathway, only T9αH-750C functions efficiently due to the absence of C13-acetylation and deacetylation capacity. Nevertheless, our results show that integration of the complete C13-acetylation–deacetylation module enables both T9αH-725A and T9αH-750C to facilitate robust baccatin III production, with T9αH-750C retaining superior catalytic performance in both pathway architectures. This finding expands our mechanistic understanding of how the C13 modification module reshapes the catalytic compatibility of upstream P450 hydroxylases.

Functional redundancy is a recurring theme across the taxane biosynthetic network, manifesting in multiple enzyme pairs: the tissue-differentiated C9α-hydroxylases T9αH-725A/T9αH-750C^12,17^, the acetyltransferases TAT/TAX19^22^, and the deacetylase isoforms T7dA/T7dA1 and T13dA1/T13dA2 characterized herein. Such redundancy likely confers evolutionary resilience to *Taxus* species under environmental fluctuations. More specifically, the coexistence of multiple deacetylase paralogs points to tissue**□**specific transcriptional regulation^42^, with distinct isoforms potentially partitioning competing pathway branches across phloem and pith tissues.

Collectively, this work unravels a previously unrecognized cryptic C13-acetylation–deacetylation module in the paclitaxel biosynthetic cascade and establishes it as a functionally significant branch within the multi**□** layered metabolic network of *Taxus*, as evidenced by the markedly enhanced baccatin III yields upon its integration into the reconstituted pathway. These findings not only enrich our mechanistic understanding of paclitaxel biosynthesis but also provide a versatile enzymatic toolkit for sustainable heterologous paclitaxel production.

## Data availability

All data necessary to interpret, verify and extend the research presented in the article are provided within the paper and in the Supplementary Information. The gene sequences of T13dA1 (accession no. PX848827), T13dA2 (accession no. PX848828) and T7dA1 (accession no. PX848829) are deposited in GenBank.

## Author contributions

C.L. and J.D. conceived this study and designed experiments. C.L. performed the bioinformatics analyses, biological and chemical experiments and *N. benthamiana* cultivation. R.C., K.X., D.C., and J.L. coordinated the experiments. X.S. helped to analyse the data. All the authors discussed the work. C.L. and J.D. wrote the manuscript.

## Acknowledgements

This work was supported by the National Natural Science Foundation of China (82404473; 82550003); the Special Research Fund for Central Universities, Peking Union Medical College (3332024065) and the China Postdoctoral Science Foundation (2026T190940; 2024M750252) and CAMS Innovation Fund for Medical Sciences (CIFMS-2025-I2M-XHJC-027).

## Methods

### Chemical and biological materials

Chemical standards were purchased from the following vendors: taxusin (TargetMol; TN6763), 9-dihydro-13-acetylbaccatin III (BioBioPha), baccatin VI (BioBioPha), baccatin III (Shanghai yuanye Bio-Technology Co., Ltd), 10-deacetylbaccatin III (Shanghai yuanye Bio-Technology Co., Ltd). 7-acetoxy-10-deacetylbaccatin III (**11**), 13-acetoxybaccatin III (**12**), 7,13-diacetoxybaccatin III (**13**) and 7-acetoxybaccatin III (**14**) were synthesized from 10-DAB by regioselective acetylation. The solvents used for extraction, analysis and preparative HPLC were purchased from Shanghai CINC High Purity Solvent Co., Ltd. All reagents used for chemical synthesis were purchased from J & K Scientific Ltd. (Beijing, China), Inno Chem Science & Technology Co., Ltd. (Beijing, China). All candidate genes and fragment amplifications were performed using KOD One^TM^ DNA polymerase (Toyobo Biotech). PCR product purifications were performed using the Trelief® DNA Gel Extraction Kit (Tsingke Biotechnology). Plasmid purifications were performed using EasyPure Plasmid MiniPrep Kit (Tsingke Biotechnology). Restriction enzymes were purchased from New England Biolabs (USA). DNA ligase 2 × pEASY Pro Assembly Mix were purchased from TransGen Biotech. Gateway® LR^TM^ and BP^TM^ Clonase II Enzyme Mix were purchase from Thermo Fisher Scientific.

### Transcriptome analysis and candidate genes retrieval

The culture methodology for inducing paclitaxel production in *Taxus × media* cell cultures, along with the corresponding metabolite profiling and transcriptome sequencing analysis, has been previously established and reported in the literature^20^. RNA sequencing and transcriptome analysis were established by Berry Genomics (Beijing, China). The chromosome-level genome of *T. chinensis* var. *mairei* was used as the reference genome. All transcripts annotated as carboxylesterase were retrieved from transcriptome database of paclitaxel-producing *T. × media* cell cultures.

### Transient expression of *Taxus* genes in *N. benthamiana*

RNA isolated from *T. × media* cell cultures was converted into cDNA using the SMARTScribe^TM^ Reverse Transcriptase (Takara). In general, the open reading frames (ORFs) of the diterpenoid boost genes (DXS and GGPPS), and taxane tailoring genes were cloned from the above cDNA. The PCR products were cloned (via Gateway cloning) into pEAQ-HT-DEST1 vectors and transferred into *Agrobacterium tumefaciens* (strain GV3101) cells using the freeze–thaw method. For routine *N. benthamiana* infiltration experiments, individual *Agrobacterium* preserved in 30% glycerol stocks were streaked out on Luria-Bertani (LB) solid agar medium containing kanamycin (50 μg ml^-1^) and rifampicin (25 μg ml^-1^) and grown for around one to two days at 30**□** °C. Patches of cells were scraped off from individual plates using sterile pipette tips and transfer them into 10 mL of LB with antibiotics (50 μg ml^-1^ kanamycin and 25 μg ml^-1^ rifampicillin) for 10 h at 30 °C and 200 rpm. then cells were centrifuged at 2,000 g for 10 min and the supernatant was removed. The cell pellet was re-suspended in 10 mL infiltration buffer (10 mM MES pH 5.6, 10 mM MgCl_2_ and 150 μM acetosyringone) and centrifuged at 2,000 g for 10 min. The supernatant was removed and the cell pellet was re-suspended in 5 mL infiltration buffer. For individually tested strains, *Agrobacterium* suspensions were diluted to an optical density at 600**□**nm (OD_600_) 0.4. When multiple constructs were infiltrated simultaneously, the corresponding *A. tumefaciens* cell cultures were mixed and the final OD_600_ of each remained 0.2. The suspensions were briefly vortexed to homogeneity and incubated at room temperature for 3 h in dark. Leaves of four-week-old *N. benthamiana* were infiltrated using needleless 1-ml syringes from the abaxial side. For experiments that require substrate injection, 4 days after *Agrobacterium* infiltration, 0.2 mM substrates in water with 1% DMSO was infiltrated into the underside side of previously *Agrobacterium*-infiltrated leaves with a needleless 1-mL syringe. Leaves were collected after 12 h post-infiltration, extracted and analyzed by HPLC and LC-MS. For experiments not requiring substrate injection, leaf samples should be harvested six days after *Agrobacterium* infiltration.

For the reconstitution of pathways that involve TBT, the following modifications were made to the procedure above to increase the production of the desired benzoylated products based on a previously reported work^12^: 1**□** mM benzoic acid was added to the induction buffer and the pH was adjusted to 5.6 before being used for the resuspension of *Agrobacterium* and preparation of the final infiltration solution.

### Truncation of target genes

The diterpene synthase (TXS) and geranylgeranyl pyrophosphate synthase (GGPPS) sequences were analyzed using the ChloroP 1.1 Prediction Server for the presence of a chloroplast transit peptide^32^. N-terminal truncation variants were designed according to the prediction algorithm’s results (Supplementary Table S2).

### Metabolite extraction and qualitative analysis of *N. benthamiana* leaves

The leaf samples were flash-frozen and lyophilized for 48 h. Methanol was used to extract metabolites from tobacco leaves. Specifically, accurately weigh 50 mg of freeze-dried powder and suspend it in 1 mL of methanol. The extraction mixture was sonicated for 30**□** min in a water bath and spun down, and the supernatant was filtered through a 0.22-*µ*m polyvinylidene difluoride filter before LC–MS analysis. LC-MS data were recorded on a Thermo Scientific Q Exactive Focus (Waltham, MA, USA). Data were collected with Thermo Xcalibur 4.0. Separation was done using a CAPCELL PAK C18 (MG III, 250 × 4.6 mm, 5 μm) with a mixture of 0.1% formic acid in water (A) and acetonitrile (B) at a constant flow rate of 1 mL per min at room temperature. The injection volume was 10**□**μL. The following gradient of solvent B was used: 0–30**□** min, 10%–70%B; 30–40**□**min, 70%–100%B; 40–50**□** min, 100%B; 50–51**□** min, 100%–10%B; 51–60**□**min 10%B. MS data were collected using electrospray ionization (ESI) in positive mode with a mass range of 100–1,500**□** *m/z*. The ionization source was set as follows: 350**□** °C Aux gas heater temperature, 35 psi Sheath gas flow rate, 10**□**arb Aux flow rate, 3,300**□** V Spray voltage, 320°C Capillary temperature. The split ratio is 1: 2. MS/MS fragmentations were generated using [M + H]^+^ as the precursor ion and fragmented with a collision energy of 15**□**eV unless otherwise stated.

### Purification of proteins and *in vitro* enzymatic assays

All proteins were purified from standard pET28a vectors expressed in *E. coli* Transetta (DE3) (TransGen Biotech, China). In detail, the pET-28a (+) plasmid was digested with Nde I and Sal I restriction enzymes (New England Biolabs). The coding sequences of target gene was amplified from the pEAQ-HT-DEST1 vectors with primers that contained homologous arms (Supplementary Table S1) and then ligated into pET-28a (+) vectors using ClonExpress® II One Step Cloning Kit (Vazyme Biotech, China). The recombinant plasmids were transformed into *E. coli* Transetta (DE3) for heterologous expression. Proteins were purified as previously described^20^, with post-lysis steps done at 4**□** °C. In brief, cells were grown to an OD_600_ of 0.6–0.8, induced with 0.3**□**mM IPTG and expressed for 18**□** h overnight at 16**□** °C. Cell pellets (For example, harvest bacterial cells from 500 mL of bacterial solution) were lysed in 25 mL lysis buffer (100 mM sodium phosphate buffer, 20 mM imidazole, pH 7.5, 1 mM PMSF) by sonication in an ice bath, clarified by centrifugation for 1 h at 8,000g. The resulting supernatant (take 2 L of bacterial solution as an example) was applied to a 1 mL column of Ni-NTA resin (GE, USA) that was pre-equilibrated with binding buffer (100 mM sodium phosphate buffer, 20 mM imidazole, pH 7.5). The resin was washed with 15 mL binding buffer and then washed with 10 mL elution buffer (100 mM sodium phosphate buffer, 50 mM imidazole, pH7.5) and 10 mL elution buffer (100 mM sodium phosphate buffer, 250 mM imidazole, pH7.5) under a flow rate of 1 mL/min at 0 °C. The purified protein was concentrated and desalted into assay buffer (50 mM PBS, 1**□**mM DTT, 10% glycerol, pH7.0) using 30 kDa ultrafiltration tubes (Millipore, USA). The protein purity was confirmed by sodium dodecyl sulfate-polyacrylamide gel electrophoresis (SDS-PAGE, Supplementary Fig. S2), and the protein concentration was determined by the Protein Quantitative Kit.

For the large-scale *in vitro* enzymatic assay of T13dA1, the reaction mixtures containing 6 mg (0.01 mM) 9-dihydro-13-acetylbaccatin III (**10**) and 2 mg of purified T13dA1 protein in a final volume of 20 mL 50 mM Tris-HCl buffer (pH 8.0) were incubated at 30°C for 5 h, and then were quenched by addition of 10 mL MeOH. The mixture was diluted with 10 mL of water and subsequently extracted with ethyl acetate (30 mL × 5). The combined organic layers were washed with brine, dried over anhydrous Na_2_SO_4_, and concentrated under reduced pressure to give crude **10a**, which was further purified by semi-preparative HPLC (eluent: 36% MeCN/H_2_O, v/v; 3 mL/min, CAPCELL PAK C18, UG80, 5 μm) to give **10a** (4 mg, approximately 68%, isolated yield). ^1^H-NMR (400 MHz, CD_3_OD and ^13^C-NMR (100 MHz, CD_3_OD) see Supplementary Table S1.

### Quantification of baccatin III

The sample preparation procedure has been described above. Separation was done using a CAPCELL PAK C18 (MG III, 250 × 4.6 mm, 5 μm) with a mixture of 0.1% formic acid in water (A) and acetonitrile (B) at a constant flow rate of 1**□** mL per min at 30**□** °C. The injection volume was 10**□**μL. The following gradient of solvent B was used: 0–30**□** min, 10%–70%B; 30–40**□**min, 70%–100%B; 40–50**□**min, 100%B; 50–51**□** min, 100%–10%B and 51–60**□** min, 10%B. The 587.24869 ion transition at a collision energy of 15**□** eV as the qualifier. The ionization source was set as follows: 350**□**°C Aux gas heater temperature, 60 psi**□** Sheath gas flow rate, 20**□** arb Aux flow rate, 3,300**□** V Spray voltage, 350 °C Capillary temperature. Quantitative analysis was performed using the Parallel Reaction Monitoring (PRM) mode. MS/MS fragmentations were generated using [M + H]^+^ as the precursor ion and fragmented with a collision energy of 15**□** eV unless otherwise stated.. The yield of baccatin III was quantitatively calculated by the standard curve method. The standard curve and regression equation of baccatin III were established as follows:

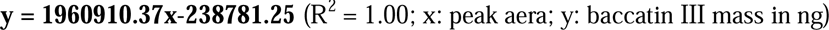

### Chemical synthesis of compounds 11−14

10-DAB (215 mg, 0.40 mmol, 1 eq., purchased from BioBioPha) was dissolved in 10 mL anhydrous dichloromethane (DCM), then Et_3_N (60.6 mg, 0.60 mmol, 1.5 eq.), DMAP (10 mg, 0.08 mmol, 0.2 eq.) and Ac_2_O (0.4 mL, 4.0 mmol, 10 eq.) was added and the mixture was stirred at room temperature under Ar atmosphere. The reaction was quenched with saturated aqueous NaHCO_3_ upon observation of three distinct product spots by thin-layer chromatography (TLC, n-Hexane: EtOAc = 1: 1, v/v), and aqueous layer was extracted with EtOAc (20 mL × 5). The combined organic phase was washed with brine, dried with anhydrous sodium sulfate, filtered and concentrated in vacuum. The resulted residue was purified on silica chromatography gel (n-Hexane: EtOAc = 1: 1) to give 7-acetoxy-10-deacetylbaccatin III (**11**, 35 mg, 15%), 7-acetoxybaccatin III (**14**, 62 mg, 25%) and 7,13-diacetoxybaccatin III (**13**, 120 mg, 45%). **13** was next enzymatically catalyzed by T7dA1 in vitro to form **12**.

**11.** HR-ESI-MS: [M + H]^+^ calc. for [C_31_H_39_O_11_]^+^ *m/z* = 587.24869; found *m/z* = 587.24884. ^1^H-NMR (CDCl_3_, 400 MHz) δ_H_8.11 (2H, d, *J* = 7.6 Hz, CH), 7.61 (1H, t, *J* = 7.6 Hz, CH), 7.48 (2H, m, CH), 5.63 (1H, d, *J* = 6.6 Hz, CH), 5.52 (1H, dd, *J* = 10.6, 7.2 Hz, CH), 4.97 (1H, dd, *J* = 9.5, 2.0 Hz, CH), 4.87 (1H, t, *J* = 7.2 Hz, CH), 4.35 (1H, d, *J* = 8.0 Hz, CH), 4.20 (1H, d, *J* = 8.4 Hz, CH), 4.09 (1H, d, *J* = 7.0 Hz, CH), 2.55 (1H, m, CH), 2.30 (3H, s, CH_3_), 2.11 (3H, br s, CH_3_), 1.99 (3H, s, CH_3_), 1.91 (1H, m, CH), 1.84 (3H, s, CH_3_), 1.08 (3H, s, CH_3_), 1.06 (3H, s, CH_3_). ^13^C NMR (100 MHz, CDCl_3_, ppm) δ_C_ 211.4 (C_q_), 170.7 (C_q_), 169.7 (C_q_), 167.0 (C_q_), 142.7 (C_q_), 134.6 (C_q_), 133.7 (CH), 130.1 (CH), 129.3 (C_q_), 128.7 (CH), 83.8 (CH), 80.4 (C_q_), 78.8 (C_q_), 76.5 (CH_2_), 75.1 (CH), 74.7 (CH), 72.0 (CH), 67.9 (CH), 56.5 (C_q_), 46.6 (CH), 42.5 (C_q_), 38.7 (CH_2_), 33.4 (CH_2_), 26.5 (CH_3_), 22.5 (CH_3_), 20.9 (CH_3_), 19.6 (CH_3_), 14.9 (CH_3_), 10.8 (CH_3_).

**12.** HR-ESI-MS: [M + H]^+^ calc. for [C_33_H_41_O_12_]^+^ *m/z* = 629.25925; found *m/z* = 629.25885. ^1^H-NMR (CD_3_OD, 400 MHz) δ_H_8.10 (2H, d, *J* = 7.6 Hz, CH), 7.64 (1H, t, *J* = 7.6 Hz, CH), 7.54 (2H, m, CH), 6.15 (1H, d, *J* = 6.6 Hz, CH), 5.73 (1H, dd, *J* = 10.6, 7.2 Hz, CH), 5.01 (1H, dd, *J* = 9.5, 2.0 Hz, CH), 4.33 (1H, d, *J* = 8.0 Hz, CH), 4.21 (1H, s, CH), 3.90 (1H, d, *J* = 7.0 Hz, CH), 2.43 (3H, s, CH_3_), 2.20 (3H, br s, CH_3_), 1.98 (3H, s, CH_3_), 1.67 (3H, s, CH_3_), 1.17 (3H, s, CH_3_). ^13^C NMR (100 MHz, CD_3_OD, ppm) δ_C_ 205.2 (C_q_), 172.1 (C_q_), 171.7 (C_q_), 167.6 (C_q_), 142.5 (C_q_), 134.9 (C_q_), 134.6 (CH), 131.1 (CH), 129.8 (C_q_), 129.7 (CH), 85.9 (CH), 82.4 (C_q_), 79.0 (C_q_), 77.5 (CH), 77.0 (CH_2_), 76.0 (CH), 72.4 (CH), 71.1 (CH), 67.9 (CH), 59.4 (C_q_), 44.5 (CH), 41.6 (C_q_), 37.6 (CH_2_), 37.1 (CH_2_), 26.9 (CH_3_), 22.9 (CH_3_), 21.2 (CH_3_), 20.8 (CH_3_), 15.0 (CH_3_), 10.4 (CH_3_).

**13.** HR-ESI-MS: [M + H]^+^ calc. for [C_35_H_43_O_13_]^+^ *m/z* = 671.26982; found *m/z* = 671.26947. ^1^H-NMR (CDCl_3_, 400 MHz) δ_H_8.08 (2H, d, *J* = 7.6 Hz, CH), 7.61 (1H, t, *J* = 7.6 Hz, CH), 7.48 (2H, m, CH), 6.25 (1H, s, CH), 6.16 (1H, t, *J* = 6.4 Hz, CH), 5.66 (1H, d, *J* = 5.6 Hz, CH), 5.58 (1H, dd, *J* = 5.6, 8.4 Hz, CH), 4.98 (1H, d, *J* = 7.6 Hz, CH), 4.33 (1H, d, *J* = 6.8 Hz, CH_2_), 4.16(1H, d, *J* = 6.8 Hz, CH_2_), 3.96 (1H, d, *J* = 5.6 Hz, CH), 2.61 (1H, m, CH_2_), 2.34 (3H, s, CH_3_), 2.25 (1H, d, *J* = 7.2 Hz, CH_2_), 2.21 (3H, s, CH_3_), 2.17 (3H, s, CH_3_), 2.03 (3H, s, CH_3_), 1.96 (3H, s, CH_3_), 1.79 (3H, s, CH_3_), 1.20 (3H, s, CH_3_), 1.16 (3H, s, CH_3_). ^13^C NMR (100 MHz, CDCl_3_, ppm) δ_C_ 202.2 (C_q_), 170.6 (C_q_), 170.4 (C_q_), 169.7 (C_q_), 169.1 (C_q_), 167.1(C_q_), 141.5 (C_q_), 133.9 (C_q_), 132.5 (CH), 130.2 (CH), 129.2 (C_q_), 128.8 (CH), 84.1 (CH), 81.0 (C_q_), 78.9 (C_q_), 76.4 (CH_2_), 75.5 (CH), 74.5 (CH), 71.5 (CH), 69.6 (CH), 56.1 (C_q_), 47.3 (CH), 43.2 (C_q_), 35.6 (CH_2_), 33.5 (CH_2_), 26.5 (CH_3_), 22.6 (CH_3_), 21.4 (CH_3_), 21.3 (CH_3_), 20.9 (CH_3_), 20.8 (CH_3_), 14.9 (CH_3_), 10.9 (CH_3_).

**14.** HR-ESI-MS: [M + H]^+^ calc. for [C_33_H_41_O_12_]^+^ *m/z* = 629.25925; found *m/z* = 629.25848. ^1^H-NMR (CDCl_3_, 400 MHz) δ_H_8.11 (2H, d, *J* = 7.6 Hz, CH), 7.61 (1H, t, *J* = 7.6 Hz, CH), 7.48 (2H, m, CH), 6.27 (1H, s, CH), 5.63 (1H, d, *J* = 7.2 Hz, CH), 5.60 (1H, m, CH), 4.99 (1H, d, *J* = 7.6 Hz, CH), 4.86 (1H, t, *J* = 6.8 Hz, CH), 4.33 (1H, d, *J* = 6.8 Hz, CH_2_), 4.16(1H, d, *J* = 6.8 Hz, CH_2_), 4.01 (1H, d, *J* = 5.6 Hz, CH), 2.63 (1H, m, CH_2_), 2.30 (3H, s, CH_3_), 2.28 (1H, d, *J* = 7.2 Hz, CH_2_), 2.18 (3H, s, CH_3_), 2.11 (3H, s, CH_3_), 2.04 (3H, s, CH_3_), 1.80 (3H, s, CH_3_), 1.14 (3H, s, CH_3_), 1.09 (3H, s, CH_3_), 1.16 (3H, s, CH_3_). ^13^C NMR (100 MHz, CDCl_3_, ppm) δ_C_ 202.4 (C_q_), 170.6 (C_q_), 170.5 (C_q_), 169.0 (C_q_), 167.0(C_q_), 144.6 (C_q_), 133.7 (C_q_), 131.6 (CH), 130.1 (CH), 129.3 (C_q_), 128.6 (CH), 84.0 (CH), 80.7 (C_q_), 78.6 (C_q_), 76.4 (CH_2_), 75.9 (CH), 74.4 (CH), 71.6 (CH), 67.9 (CH), 56.2 (C_q_), 47.5 (CH), 42.8 (C_q_), 38.4 (CH_2_), 33.4 (CH_2_), 26.7 (CH_3_), 22.6 (CH_3_), 21.1 (CH_3_), 20.8 (CH_3_), 20.1 (CH_3_), 15.2 (CH_3_), 10.7 (CH_3_).

### NMR analysis

All the NMR data were recorded on BRUKER-AVANCEIII-400 (CADM-YQ-003) in the CD_3_OD and CDCl_3_ and the data was processed and visualized on MestRenova (version 14.0.0-23239). The chemical shift (δ) of ^1^H NMR was given in ppm relative to Me_4_Si (TMS) (δ = 0.00 ppm) and CD_3_OD (δ = 3.30 ppm). The chemical shift (δ) of ^13^C NMR was given in ppm relative to CD_3_OD (δ = 49.00 ppm) and TMS (δ = 0.00 ppm).

## References

1. Tang, X. et al. Versatile 2□oxoglutarate-dependent dioxygenases catalyze radical-mediated multifunctional skeleton reconstructions and oxidation modifications of taxoids. J. Am. Chem. Soc. 148, 11333–11343 (2026).

2. Xiong, X. et al. The *Taxus* genome provides insights into paclitaxel biosynthesis. Nat. Plants 7, 1026–1036 (2021).

3. Liang, F. et al. Elucidation of the final steps in Taxol biosynthesis and its biotechnological production. Nat. Syn. 4, 1212–1222 (2025).

4. Borah, J. C., Boruwa, J. & Barua, N. C. Synthesis of the C-13 side-chain of Taxol. Curr. Org. Synth. 4, 175–199 (2007).

5. Sharma, A. et al. An overview on Taxol production technology and its applications as anticancer agent. Biotechnol. Bioproc. E. 27, 706–728 (2022).

6. Sabzehzari, M., Zeinali, M. & Naghavi, M. R. Alternative sources and metabolic engineering of Taxol: Advances and future perspectives. Biotechnol. Adv. 43, 107569 (2020).

7. Tan, C. et al. A synthetic biology roadmap for sustainable production of the plant-originated anti-cancer drug paclitaxel. Trends Biotechnol. 44, 1939–1959 (2026).

8. Chen, Q. et al. Recent advances and perspectives in biosynthesis of paclitaxel: key enzymes and intermediates. Int. J. Biol. Macromol. 331, 148049 (2025).

9. Lai, M. et al. Designing an oxidase toolbox for site-directed oxidation of taxanes. Nat. Commun. 17, 841 (2026).

10. Jiang, B. et al. Characterization and heterologous reconstitution of *Taxus* biosynthetic enzymes leading to baccatin III. Science 383, 622–629 (2024).

11. Wildung, M. R. & Croteau, R. A cDNA clone for taxadiene synthase, the diterpene cyclase that catalyzes the committed step of Taxol biosynthesis. J. Biol. Chem. 271, 9201–9204 (1996).

12. McClune, C. J. et al. Discovery of FoTO1 and Taxol genes enables biosynthesis of baccatin III. Nature 643, 582–592 (2025).

13. Croteau, R., et al. Taxol biosynthesis and molecular genetics. Phytochem. Rev. 5, 75–97 (2006).

14. Jennewein, S., Long. R. M., Williams, R. M. & Croteau, R. Cytochrome P450 taxadiene 5α-hydroxylase, a mechanistically unusual monooxygenase catalyzing the first oxygenation step of Taxol biosynthesis. Chem. Biol. 11, 379–387 (2004).

15. Jennewein, S., Rithner, C. D., Williams, R. M. & Croteau, R. B. Taxol biosynthesis: taxane 13α-hydroxylase is a cytochrome P450-dependent monooxygenase. Proc. Natl Acad. Sci. USA 98, 13595–13600 (2001).

16. Schoendorf, A., Rithner, C. D., Williams, R. M. & Croteau, R. B. Molecular cloning of a cytochrome P450 taxane 10β-hydroxylase cDNA from *Taxus* and functional expression in yeast. Proc. Natl Acad. Sci. USA 98, 1501–1506 (2001).

17. Zhang, Y. et al. Synthetic biology identifies the minimal gene set required for paclitaxel biosynthesis in a plant chassis. Mol. Plant 16, 1951–1961 (2023).

18. Chau, M. D., Jennewein, S., Walker, K. & Croteau, R. Taxol biosynthesis: molecular cloning and characterization of a cytochrome P450 taxoid 7β-hydroxylase. Chem. Biol. 11, 663–672 (2004).

19. Chau, M. D. & Croteau, R. Molecular cloning and characterization of a cytochrome P450 taxoid 2α-hydroxylase involved in Taxol biosynthesis. Arch. Biochem. Biophys. 427, 48–57 (2004).

20. Li, C. et al. Novel α-KG/Fe(II)-dependent dioxygenases catalyzing C1β-hydroxylation and construction of 5/7/6-skeleton of highly oxygenated taxoids. Angew. Chem. Int. Ed. 64, e202517041 (2025).

21. Li, C. et al. A cytochrome P450 enzyme catalyses oxetane ring formation in paclitaxel biosynthesis. Angew. Chem. Int. Ed. 63, e202407070 (2024).

22. Yang, C. et al. Biosynthesis of the highly oxygenated tetracyclic core skeleton of Taxol. Nat. Commun. 15, 2339 (2024).

23. Chau, M., Walker, K., Long, R. & Croteau, R. Regioselectivity of taxoid-*O*-acetyltransferases: heterologous expression and characterization of a new taxadien-5α-ol-*O*-acetyltransferase. Arch. Biochem. Biophys. 430, 237–246 (2004).

24. Walker, K. & Croteau, R. Molecular cloning of a 10-deacetylbaccatin III-10-*O*-acetyl transferase cDNA from *Taxus* and functional expression in *Escherichia coli*. Proc. Natl Acad. Sci. USA 97, 583–587 (2000).

25. Walker, K. & Croteau, R. Taxol biosynthesis: molecular cloning of a benzoyl-CoA: taxane 2α-*O*-benzoyltransferase cDNA from *Taxus* and functional expression in *Escherichia coli*. Proc. Natl Acad. Sci. USA 97, 13591–13596 (2000).

26. Li, C., Yin, X. & Dai, J. The history of studies on oxetane ring formation in paclitaxel biosynthesis. ChemBioChem 26, e202400947 (2025).

27. Wang, Y. et al. Natural taxanes: developments since 1828. Chem. Rev. 111, 7652–7709 (2011).

28. Hong, B., et al. Biosynthesis of strychnine. Nature 607, 617–622 (2022).

29. Caputi, L. et al. Missing enzymes in the biosynthesis of the anticancer drug vinblastine in Madagascar periwinkle. Science 360, 1235–1239 (2018).

30. Dang, T. T., Chen, X. & Facchini, P. Acetylation serves as a protective group in noscapine biosynthesis in opium poppy. Nat. Chem. Biol. 11, 104–106 (2015).

31. Peña, R. D. L. et al. Complex scaffold remodeling in plant triterpene biosynthesis. Science 379, 361–368 (2023).

32. Peña, R.D.L. and Sattely, E.S. Rerouting plant terpene biosynthesis enables momilactone pathway elucidation. Nat. Chem. Biol. 17, 205–212 (2021).

33. Eisenreich, W., Menhard, B., Hylands, P. J., Zenk, M. H. & Bacher, A. Studies on the biosynthesis of taxol: the taxane carbon skeleton is not of mevalonoid origin. Proc. Natl Acad. Sci. USA 93, 6431–6436 (1996).

34. Jennewein, S. & Croteau, R. Taxol: biosynthesis, molecular genetics, and biotechnological applications. Appl. Microbiol. Biotechnol. 57, 13–19 (2001).

35. Reed, J. & Osbourn, A. Engineering terpenoid production through transient expression in *Nicotiana benthamiana*. Plant Cell Rep. 37, 1431–1441 (2018).

36. Estevez, J. M. et al. 1-deoxy-D-xylulose-5-phosphate synthase, a limiting enzyme for plastidic isoprenoid biosynthesis in plants. J. Biol. Chem. 276, 22901–22909 (2001).

37. Forestier, E. C. F. et al. Developing a *Nicotiana benthamiana* transgenic platform for high-value diterpene production and candidate gene evaluation. Plant Biotechnol. J. 19, 1614–1623 (2021).

38. Eljounaidi, K. et al. Discovery and characterisation of terpenoid biosynthesis enzymes from *Daphniphyllum macropodum*. BMC Plant Biol. 25, 483 (2025).

39. Brückner, K. & Tissier, A. High-level diterpene production by transient expression in *Nicotiana benthamiana*. Plant Methods 9, 46 (2013).

40. Srividya, N. et al. Biochemical characterization of acyl activating enzymes for side chain moieties of Taxol and its analogs. J. Biol. Chem. 295, 4963–4973 (2020).

41. Learned, R. M., & Fink, G. R. 3-Hydroxy-3-methylglutaryl-coenzyme A reductase from *Arabidopsis thaliana* is structurally distinct from the yeast and animal enzymes. Proc. Natl Acad. Sci. USA 86, 2779–2783 (1989).

42. Yu, C. et al. Tissue-specific study across the stem of *Taxus media* identifies a phloem-specific TmMYB3 involved in the transcriptional regulation of paclitaxel biosynthesis. Plant J. 103, 95–110 (2020).

