## Supplementary Information for "Missing taxane C13α-*O*-deacetylases reveal a cryptic acetylation–deacetylation module in paclitaxel biosynthesis"

### Supplementary Tables

**Table S1**. Primers used in this study (The underlined part is homologous sequences).

| Name | Sequence 5′ to 3′ |
|---|---|
| TXS-F | GGGGACAAGTTTGTACAAAAAAGCAGGCTGCATGGCTCAGCTCTCATTTAATGC |
| TXS-R | GGGGACCACTTTGTACAAGAAAGCTGGGTGTCATACTTGAATTGGATCAATATAAACTTT |
| FoTO1-F | GGGGACAAGTTTGTACAAAAAAGCAGGCTGCATGGCAGAAACAATGAATGAGAAA |
| FoTO1-R | GGGGACCACTTTGTACAAGAAAGCTGGGTGTCATGGAGGAGTTGGATCCTTT |
| T5*α*H-F | GGGGACAAGTTTGTACAAAAAAGCAGGCTGCATGGACGCCCTGTATAAGAGC |
| T5*α*H-R | GGGGACCACTTTGTACAAGAAAGCTGGGTGCTATGGTCTCGGAAACAGTTTAATG |
| T7*β*H-F | GGGGACAAGTTTGTACAAAAAAGCAGGCTGCATGGATGCCCTTTCTCTTGTAAAC |
| T7*β*H-R | GGGGACCACTTTGTACAAGAAAGCTGGGTGTCAGGATCTGGCGATAAGTTTTAT |
| DBAT-F | GGGGACAAGTTTGTACAAAAAAGCAGGCTGCATGGCAGGCTCAACAGAATTTG |
| DBAT-R | GGGGACCACTTTGTACAAGAAAGCTGGGTGTCAAGGTTTAGTTACATATTTGTTTGTCA |
| T9*α*H (725A)-F | GGGGACAAGTTTGTACAAAAAAGCAGGCTGCATGGATTCCTTAAGTTTTCTAAAAAGC |
| T9*α*H (725A)-R | GGGGACCACTTTGTACAAGAAAGCTGGGTGTTAGGTTCTGGGAAATAGTTTAATG |
| T9*α*H (750C)-F | GGGGACAAGTTTGTACAAAAAAGCAGGCTGCATGGCATTTTCTAGACTGATTGCA |
| T9*α*H (750C)-R | GGGGACCACTTTGTACAAGAAAGCTGGGTGTTAGATGCAATTTAACAATTTCAGGGT |
| T10*β*H-F | GGGGACAAGTTTGTACAAAAAAGCAGGCTGCATGGATAGCTTCATTTTTCTGAGAA |
| T10*β*H-R | GGGGACCACTTTGTACAAGAAAGCTGGGTGTTAGGATCTCGGAAAAAGTTTTATG |
| TAT-F | GGGGACAAGTTTGTACAAAAAAGCAGGCTGCATGGAGAAGACAGATTTACATGTAAATC |
| TAT-R | GGGGACCACTTTGTACAAGAAAGCTGGGTGTCATACTTTAGCCACATATTTTTTCATC |
| TBT-F | GGGGACAAGTTTGTACAAAAAAGCAGGCTGCATGGGCAAGTTCCATGTAGATATG |
| TBT-R | GGGGACCACTTTGTACAAGAAAGCTGGGTGTTATAACTTAGAGTTACATATTTTAGCCA |
| T13*α*H-F | GGGGACAAGTTTGTACAAAAAAGCAGGCTGCATGGATGCCCTTAAGCAATTG |
| T13*α*H-R | GGGGACCACTTTGTACAAGAAAGCTGGGTGTTAAGATCTGGAATAGAGTTTAATGGG |
| T2*α*H-F | GGGGACAAGTTTGTACAAAAAAGCAGGCTGCATGGACGCCATGGATCTCAC |
| T2*α*H-R | GGGGACCACTTTGTACAAGAAAGCTGGGTGTTAGGATCGAGAAATAAGTTTAATAGGA |
| T1*β*H-F | GGGGACAAGTTTGTACAAAAAAGCAGGCTGCATGGCTTCCTCGGCGCA |
| T1*β*H-R | GGGGACCACTTTGTACAAGAAAGCTGGGTGTCAATAAATAGGGGATATGCCG |
| TOT-F | GGGGACAAGTTTGTACAAAAAAGCAGGCTGCATGGTTCATGTGTTGCAGGTAGTG |
| TOT-R | GGGGACCACTTTGTACAAGAAAGCTGGGTGTTAGGATCTGGGAGTAGGTTTTATTG |
| T13dA1-F | GGGGACAAGTTTGTACAAAAAAGCAGGCTGCATGTCCGAAGCCGAGGCT |
| T13dA1-R | GGGGACCACTTTGTACAAGAAAGCTGGGTGCTAGCTTTGGAATGAACCCATATTT |
| T13dA2-F | GGGGACAAGTTTGTACAAAAAAGCAGGCTGCATGTCGGATAAACGCGAGG |
| T13dA2-R | GGGGACCACTTTGTACAAGAAAGCTGGGTGTTAGTATTTCATTTTGACGAATTCCG |
| T7dA1-F | GGGGACAAGTTTGTACAAAAAAGCAGGCTGCATGGCGGATAAGAGCGAGC |
| T7dA1-R | GGGGACCACTTTGTACAAGAAAGCTGGGTGTTAATTAGTTAAGTTAATGAAGTGAGAGATGC |

| | |
|---|---|
| T9dA-F | GGGGACAAGTTTGTACAAAAAAGCAGGCTGCATGGAAGCAGCTATGGCGG |
| T9dA-R | GGGGACCACTTTGTACAAGAAAGCTGGGTGCTAACTTTTACTCCAGACGAAATCAGA |
| T9Ox-F | GGGGACAAGTTTGTACAAAAAAGCAGGCTGCATGGATTCCTCGCTGGAGAAA |
| T9Ox-R | GGGGACCACTTTGTACAAGAAAGCTGGGTGTCAATAAGTAGGAATTATGCCTGCAT |
| TAX19-F | GGGGACAAGTTTGTACAAAAAAGCAGGCTGCATGGAGAATGCAGTGTGGAAAG |
| TAX19-R | GGGGACCACTTTGTACAAGAAAGCTGGGTGTCATACTGCAGTCACATATTTGTTTAT |
| T7AT-F | GGGGACAAGTTTGTACAAAAAAGCAGGCTGCATGGAGAATCCAAGCTCAACAGA |
| T7AT-R | GGGGACCACTTTGTACAAGAAAGCTGGGTGTCACGATTTAGTCACAAATTTGTTT |
| T13dA1-F | gtgccgcgcggcagccatATGTCCGAAGCCGAGGCT |
| T13dA1-R | ggccgcaagcttgtcgaCTAGCTTTGGAATGAACCCATATT |
| T7dA1-F | gtgccgcgcggcagccatATGGCGGATAAGAGCGAGC |
| T7dA1-R | ggccgcaagcttgtcgaTTAATTAGTTAAGTTAATGAAGTGAGAGAT |
| tTXS | GGGGACAAGTTTGTACAAAAAAGCAGGCTGCATGGTAATGATGAGCAGCAGCAC |
| tTXS | GGGGACCACTTTGTACAAGAAAGCTGGGTGTCATACTTGAATTGGATCAATATAAACTTT |
| tGGPPS | GGGGACAAGTTTGTACAAAAAAGCAGGCTGCATGGCTTCCTATCAAGAATGCAA |
| tGGPPS | GGGGACCACTTTGTACAAGAAAGCTGGGTGTCAGTTTTGCCTGAATGCAATG |
| HMGR | GGGGACAAGTTTGTACAAAAAAGCAGGCTGCATGAAGAAAAAGCAAGCTGGTCC |
| HMGR | GGGGACCACTTTGTACAAGAAAGCTGGGTGTCATGTTGTTGTTGTTGTCGTTGT |

**Table S2:** Chloroplast transit peptide predictions using ChloroP.

| Genes | Chloroplast transit peptide residues |
|---|---|
| TXS | 1-58 |
| GGPPS | 1-18 |

**Table S3.** $^{1}$H- and $^{13}$C-NMR data of **10a** ($\delta$ in ppm, $J$ in Hz), multiplicity: s = singlet, d = doublet, m = multiplet.

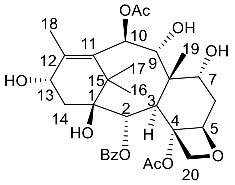

**10a**

| No. | 10a | | |
|---|---|---|---|
| | $^{1}$H | $^{13}$C | type |
| **1** | | 79.2 | $C_q$ |
| **2** | 5.73 (d, 6.0) | 75.0 | CH |
| **3** | 3.14 (d, 6.0) | 48.4 | CH |
| **4** | | 82.9 | $C_q$ |
| **5** | 4.98 (d, 8.0) | 85.6 | CH |
| **6** | 2.46 (m, $H_{\alpha}$) | 38.6 | $CH_2$ |
| | 1.81 (m, $H_{\beta}$) | | |
| **7** | 4.44 m | 75.0 | CH |
| **8** | | 45.6 | $C_q$ |
| **9** | 4.48 (d, 10.8) | 78.1 | CH |
| **10** | 6.24 (d, 10.8) | 75.0 | CH |
| **11** | | 134.9 | $C_q$ |
| **12** | | 145.3 | $C_q$ |
| **13** | 4.75 m | 68.6 | CH |
| **14** | 2.26 (m, $H_{\alpha}$) | 40.3 | $CH_2$ |
| | 2.32 (m, $H_{\beta}$) | | |
| **15** | | 43.9 | $C_q$ |
| **16** | 1.61 (s) | 23.1 | $CH_3$ |
| **17** | 1.11 (s) | 28.8 | $CH_3$ |
| **18** | 2.04 (br s) | 15.6 | $CH_3$ |
| **19** | 1.76 (s) | 13.1 | $CH_3$ |
| **20** | 4.20 (d, 8.0) | 77.6 | $CH_2$ |
| | 4.15 (d, 8.0) | | |
| **2-OBz** | 8.13 (2H. dd, 7.2, 2.0), 7.63 (1H, t, 7.2), 7.52 (2H, d, 7.2) | 167.7 | $C_q$ |
| **2-OAc** | | 134.5 | CH |
| | | 131.5*2 | |
| | | 131.2*2 | |
| | | 129.6 | |
| **4-OAc** | 2.21 (3H, s) | 172.0 | $C_q$ |
| | | 23.2 | $CH_3$ |
| **10-OAc** | 2.10 (3H, s) | 172.3 | $C_q$ |
| | | 21.3 | $CH_3$ |

$^{1}$H- (400 MHz) and $^{13}$C-NMR (100 MHz) data were recorded in $CD_3OD$.

### Supplementary Figures

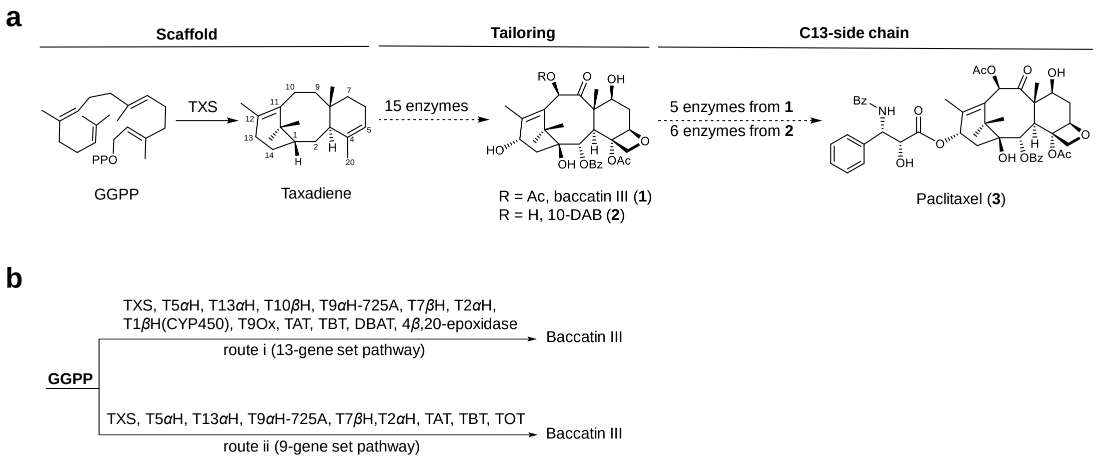

**Fig. S1. Summary of paclitaxel biosynthesis and two different cases of heterologous reconstitution of baccatin III pathways in *Nicotiana benthamiana* leaves**. **a**, Paclitaxel biosynthesis proceeds through three well-defined biosynthetic stages: (i) diterpene scaffold formation, (ii) oxidative and acylative post-modifications of the core structure, and (iii) C13 side-chain assembly and coupling. **b**, Two reported cases of heterologous reconstruction of baccatin III in tobacco lacking key genes. Route i lacks the key enzyme that mediates the oxetane ring formation and route ii lacks T10$\beta$H, T1$\beta$H and T9Ox.

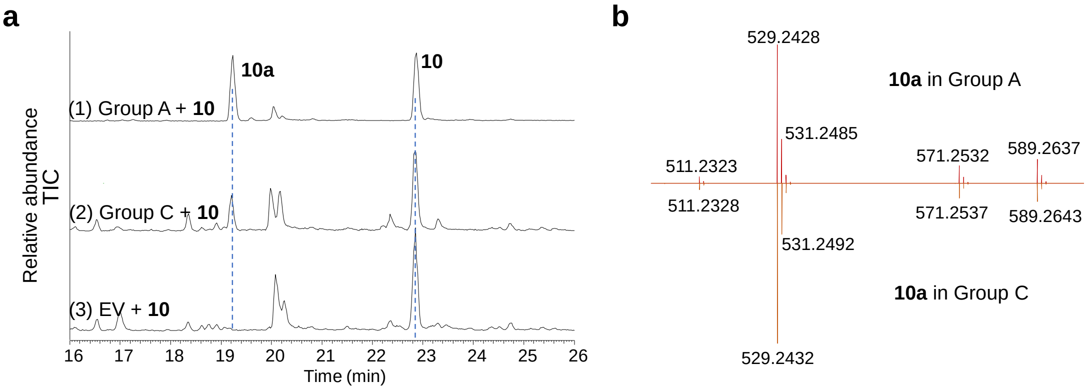

**Fig. S2. A comparative analysis of the catalytic activities when feeding 10 to *N. benthamiana* leaves expressing group A and group B. a**, Total Ion Chromatogram (TIC) of extracts from *N. benthamiana* leaves with transient expression of (group A + **10**), (group B + **10**) and empty vector (EV) + **10**. **b**, MS/MS fragmentation of **10a** produced in tobacco expressing group A and group B, respectively.

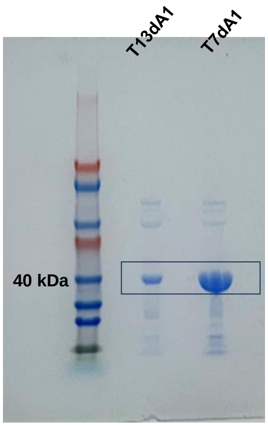

**Fig. S3. SDS-PAGE analysis of recombinant T13dA1 (cbs 52) and T7dA1 (cbs 17) purified by affinity chromatography.** predicted M.W., T13dA1: 38.28 kDa; T7dA1: 37.79 kDa.

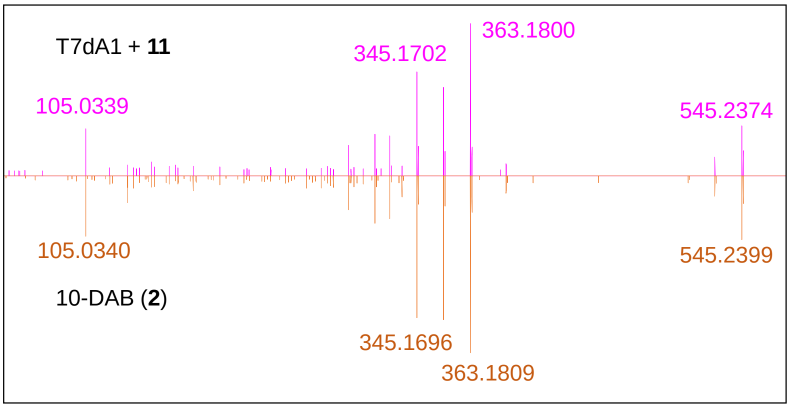

**Fig. S4. MS/MS spectra of 10-DAB (2) produced in tobacco and its standard.** MS/MS fragmentation of **2** produced in tobacco expressing T7dA1 + **11** compared to the fragmentation results of 10-DAB standard.

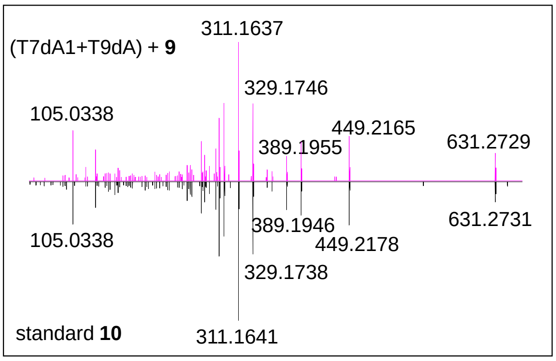

**Fig. S5. MS/MS spectra of 10 produced in tobacco and its standard.** MS/MS fragmentation of **10** produced in tobacco expressing T7dA1 + T9dA compared to the fragmentation results of the **10** standard.

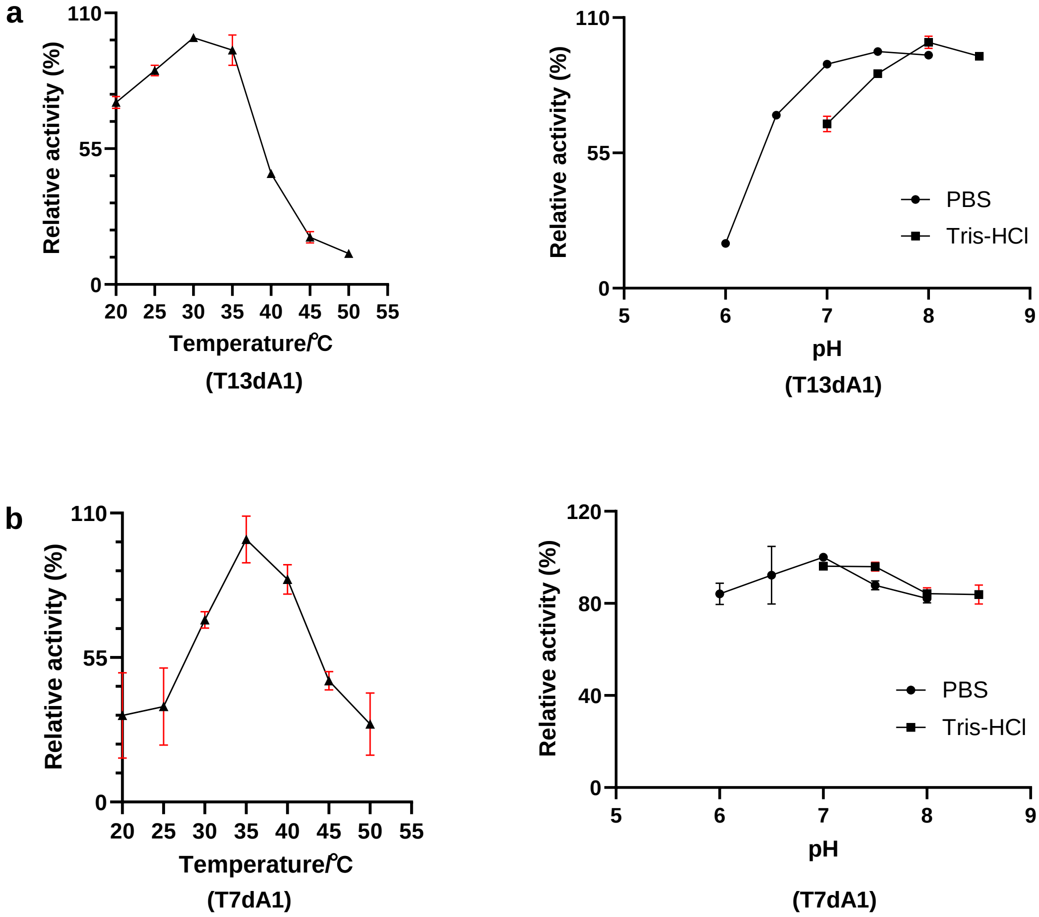

**Fig. S6. Biochemical properties of T13dA1 and T7dA1. a**, Effects of temperature and pH on T13dA1 activity toward **10** with 8 $\mu$g enzyme incubated at 30 °C for 10 min in a 100 $\mu$L buffer. **b**, Effects of temperature and pH on T7dA1 activity toward **12** with 25 $\mu$g enzyme incubated at 35 °C for 8 min in a 100 $\mu$L buffer. Data are presented as means ± standard deviation of three independent replicates ($n$ = 3). Error bars represent standard deviations.

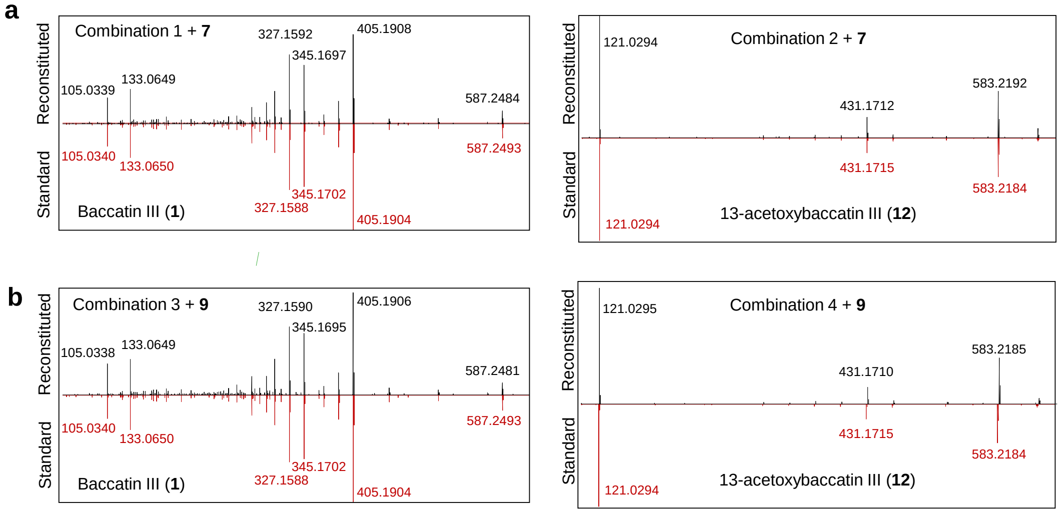

**Fig. S7. MS/MS spectra of 1 produced in tobacco and its standard.** MS/MS fragmentation of **1** produced in tobacco expressing (**a**) combination 1 + **7**, and (**b**) combination 3 + **9**, respectively compared to the fragmentation results of the **1** standard. By contrast, no baccatin III (**1**) was observed in conmbinations 2 and 4 that lacked T13dA1, whereas 13-acetoxybaccatin III (**12**) accumulated in these two control groups instead.

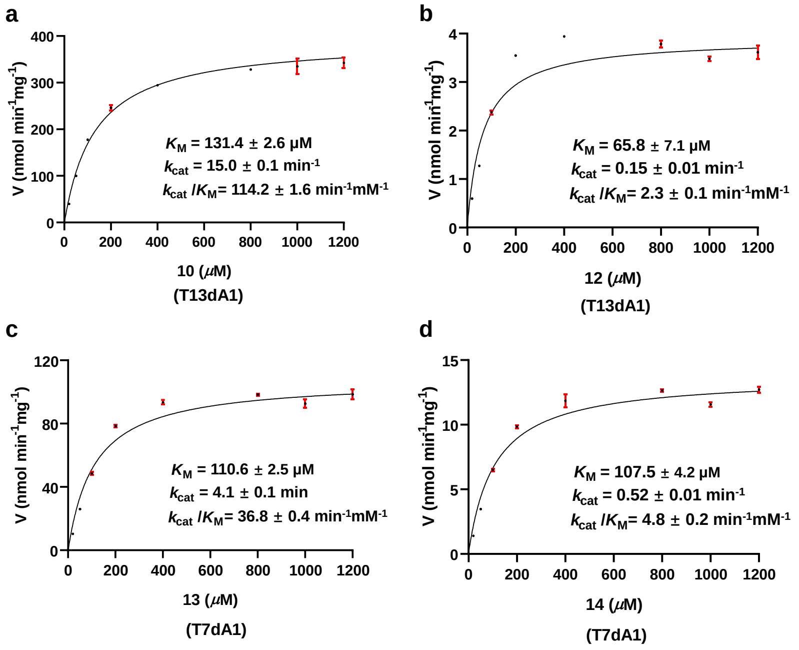

**Fig. S8. Steady-state kinetics of T13dA1 for 10 and 12 and T7dA1 for 13 and 14**. **a** and **b**, Steady-state kinetics of T13dA1 for **10** and **12** with 5 $\mu$g enzyme incubated at 30°C for 10 min in 100 $\mu$l Tris-HCl buffer (pH 8.0). **c** and **d**, Steady-state kinetics of T7dA1 for **13** and **14** with 24 $\mu$g enzyme incubated at 35°C for 8 min and 60 min respectively, in 100 $\mu$l PBS buffer (pH 8.0). Data are presented as means ± standard deviation of three independent replicates ($n$ = 3). Error bars represent standard deviations.

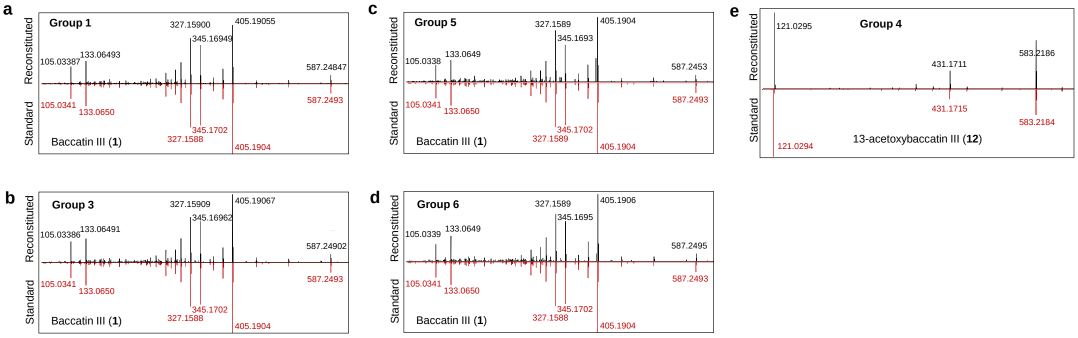

**Fig. S9.** MS/MS spectra of baccatin III (**1**) and 13-acetoxybaccatin III (**12**) produced in tobacco compared to their standards. For baccatin III, the MS/MS fragmentation of **1** produced in tobacco expressing **a**, Group 1, **b**, Group 3, **c**, Group 5, and **d**, Group 6 are shown respectively against the fragmentation results of the **1** standard. In addition, the MS/MS fragmentation of **12** produced in tobacco expressing **e**, Group 4 with its standard.

Thermo Qexactive Focus Report

compound NO. : 20-3-2
Method : LCMS(compound)-low

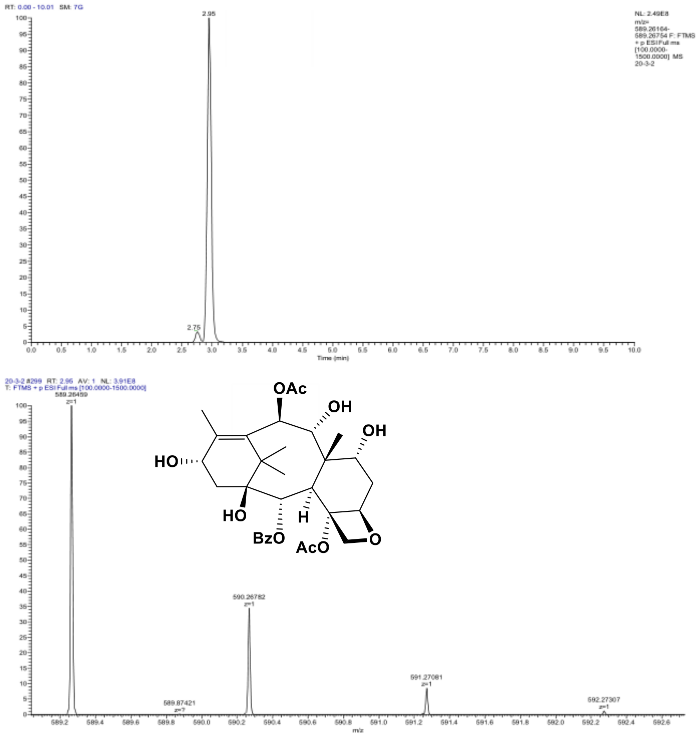

| m/z | Theo. Mass | Delta (ppm) | RDB equiv. | Composition | |
|---|---|---|---|---|---|
| 589.26459 | 589.26434 | 0.43 | 11.5 | C31 H41 O11 | M+H |

**Fig. S10. HR-ESI-MS spectrum of 10a.**

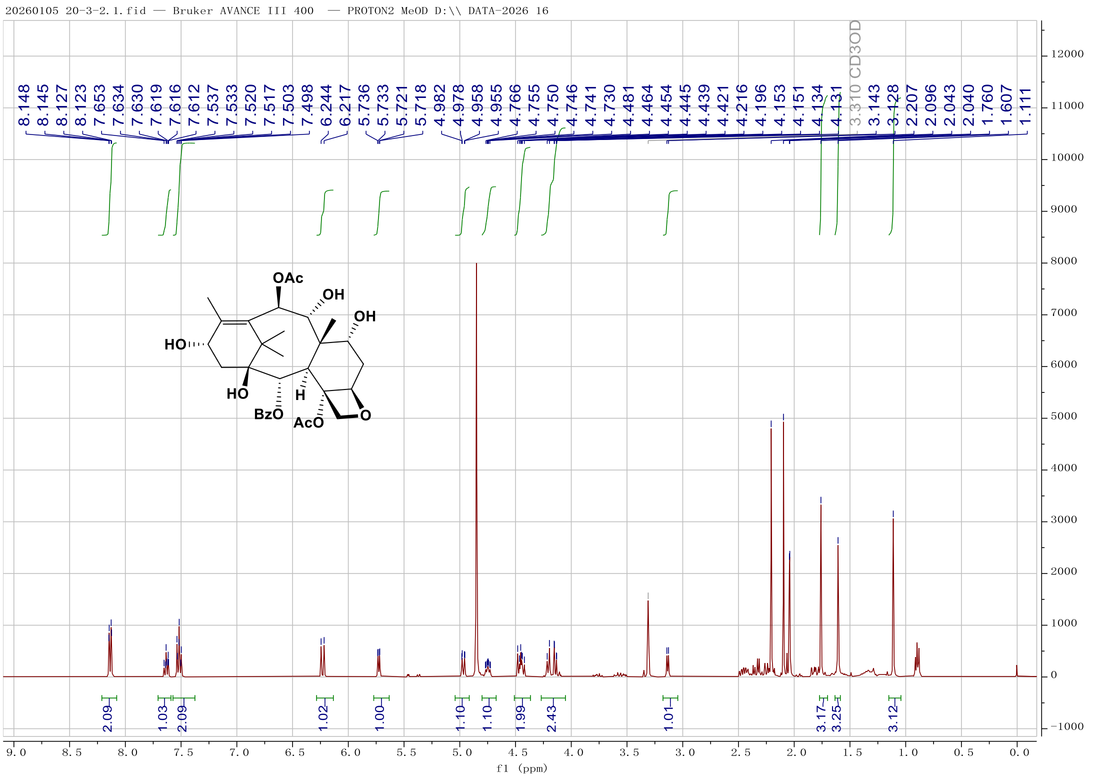

**Fig. S11. [1]H-NMR spectrum of 10a ($CD_3OD$, 400 MHz).**

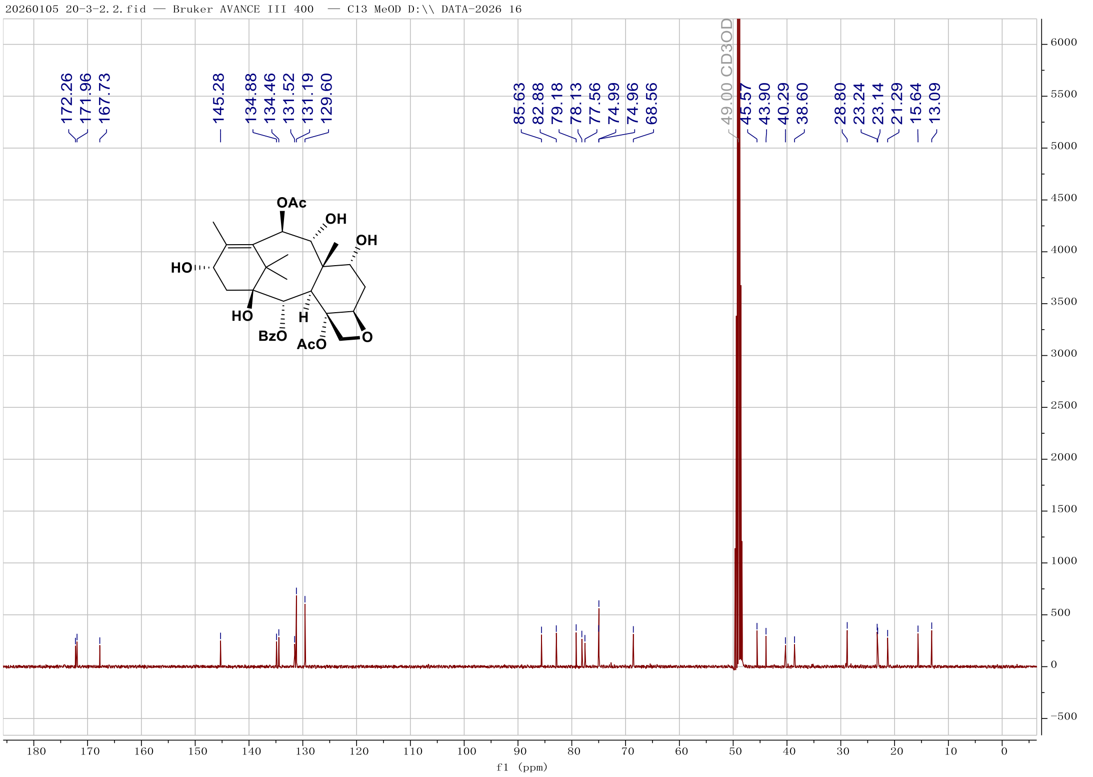

**Fig. S12. [13]C-NMR spectrum of 10a ($CD_3OD$, 100 MHz).**

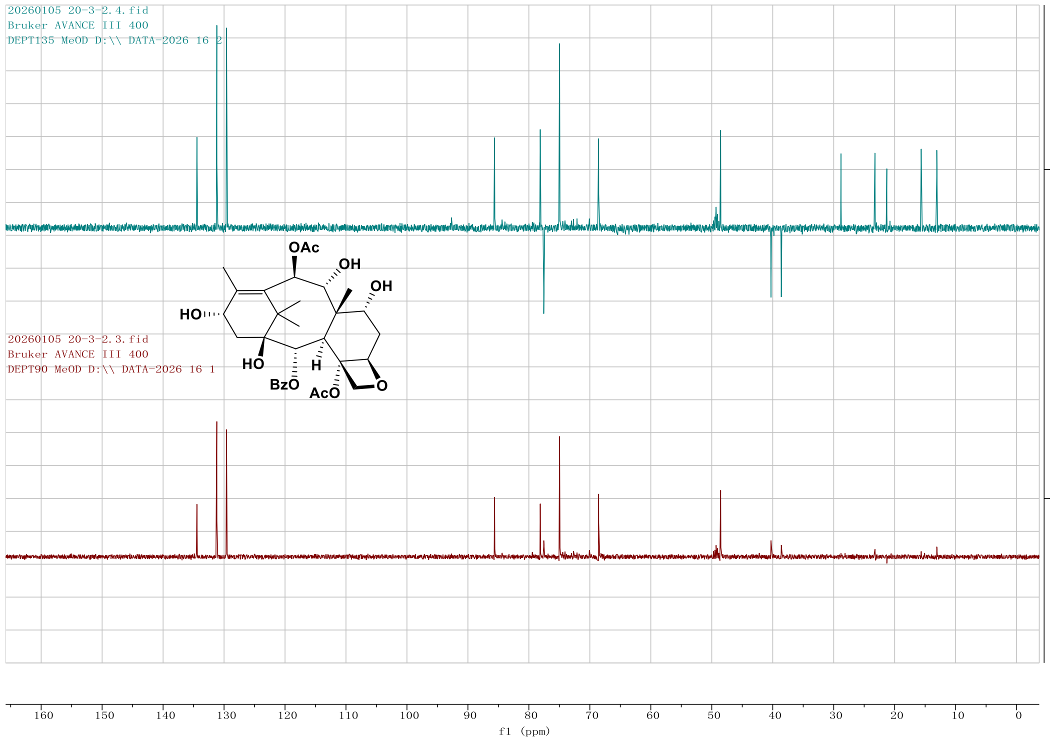

**Fig. S13. DEPT spectrum of 10a in $CD_3OD$.**

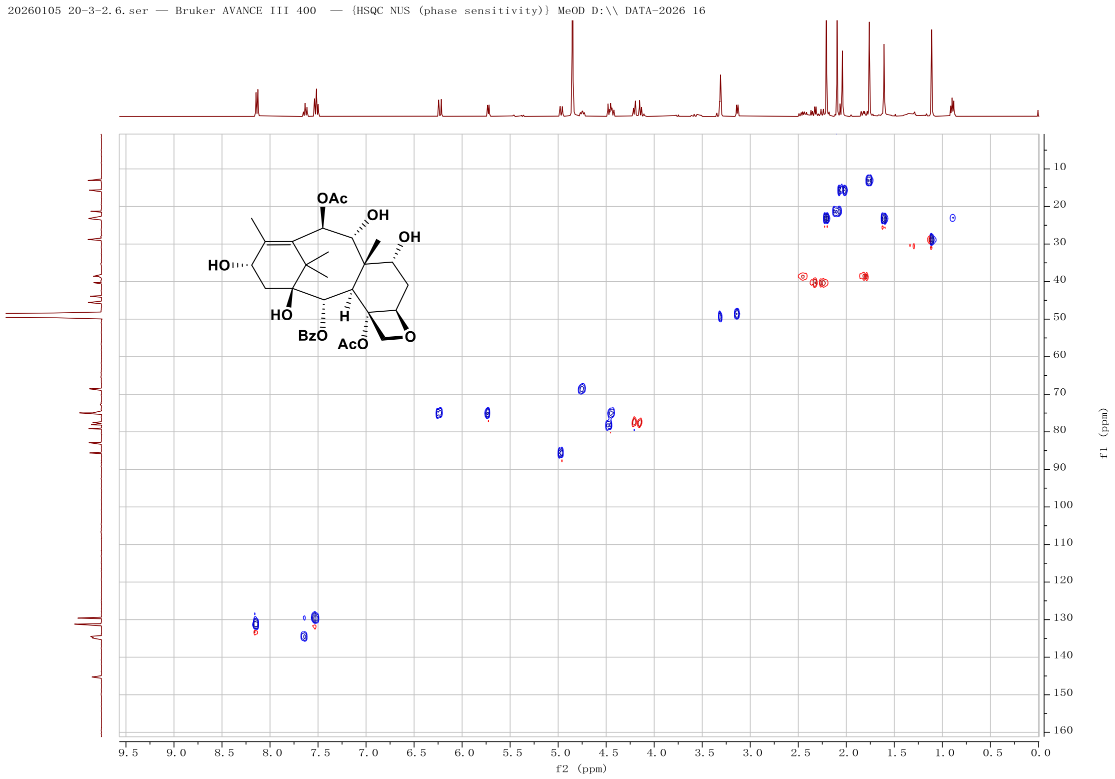

**Fig. S14. HSQC spectrum of 10a in $CD_3OD$.**

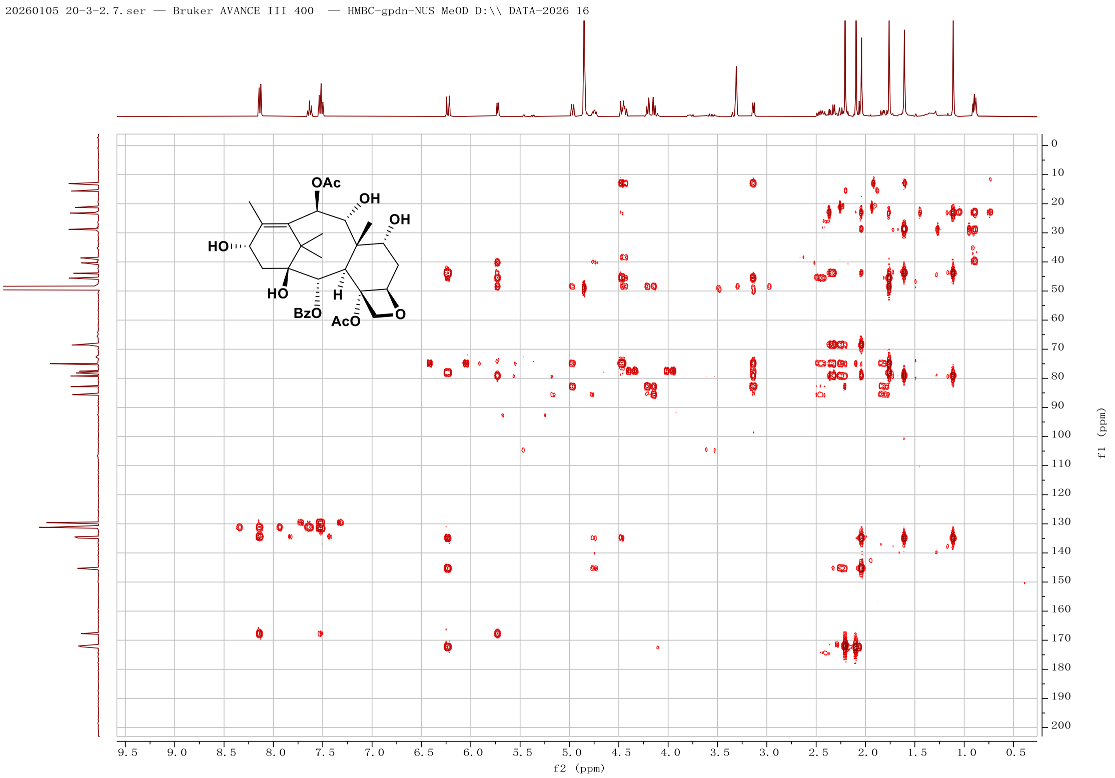

**Fig. S15. HMBC spectrum of 10a in $CD_3OD$.**

Thermo Qexactive Focus Report

compound NO. : F2-ZF

Method : LCMS(compound)-low

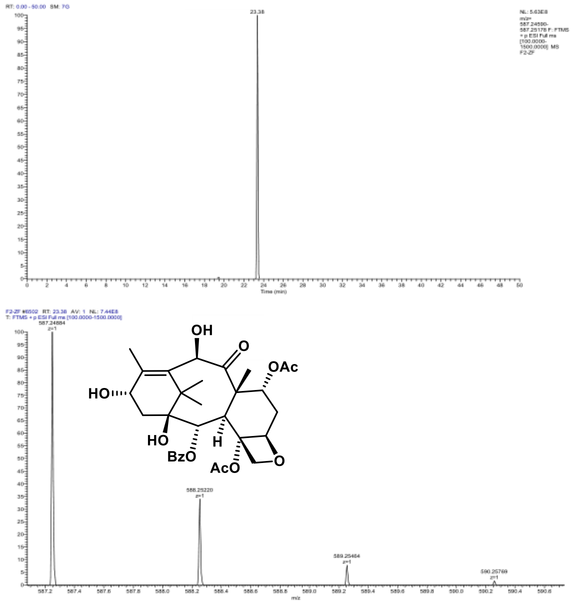

| m/z | Theo. Mass | Delta (ppm) | RDB equiv. | Composition | |
|---|---|---|---|---|---|
| 587.24884 | 587.24869 | 0.26 | 12.5 | C31 H39 O11 | M+H |

**Fig. S16. HR-ESI-MS spectrum of 11.**

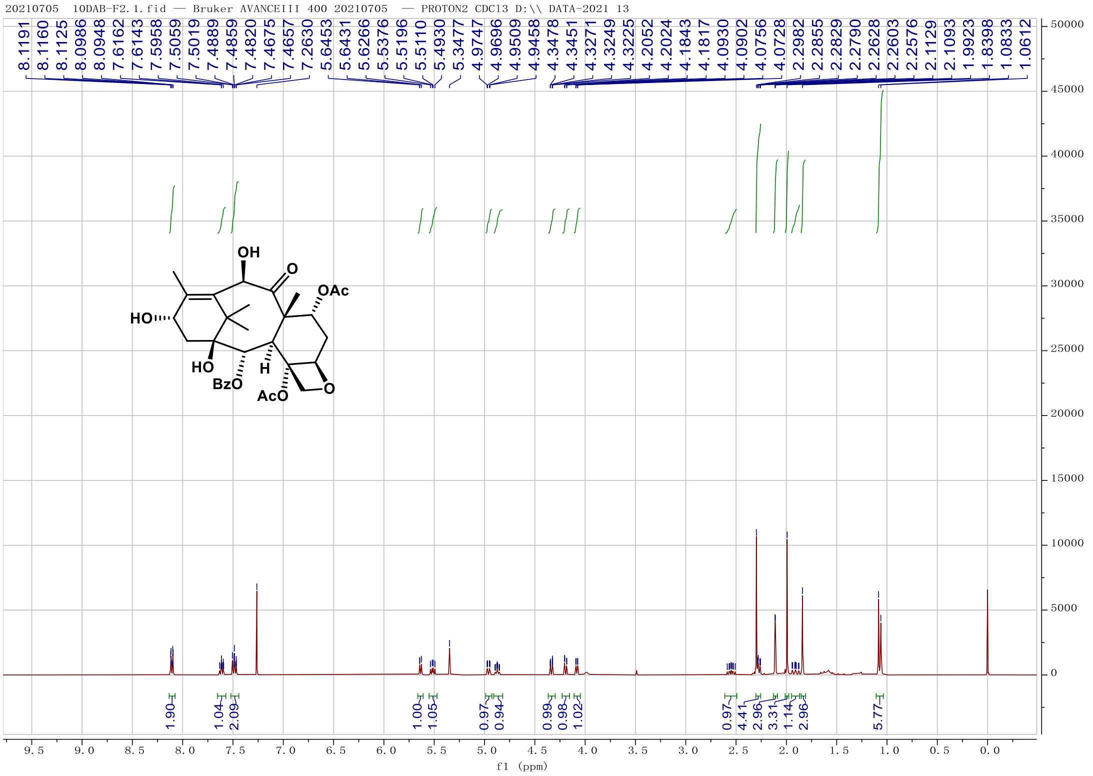

**Fig. S17. $^{1}$H-NMR spectrum of 11 ($CDCl_3$, 400 MHz).**

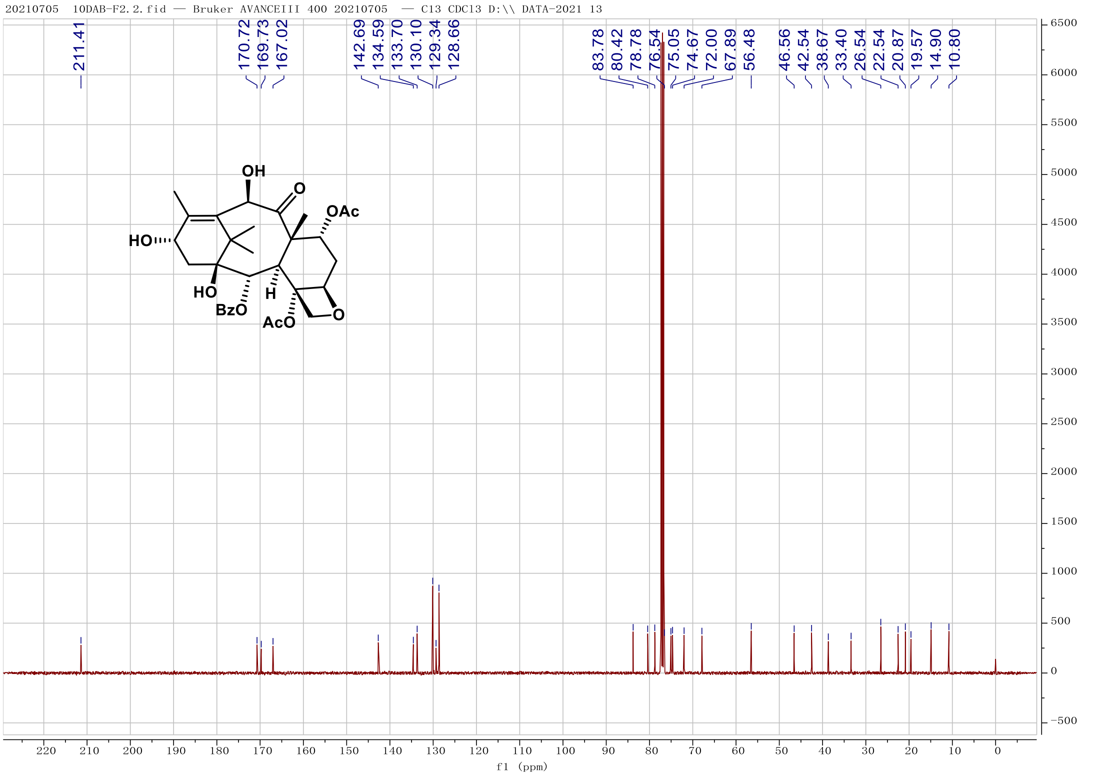

**Fig. S18. $^{13}$C-NMR spectrum of 11 ($CDCl_3$, 100 MHz).**

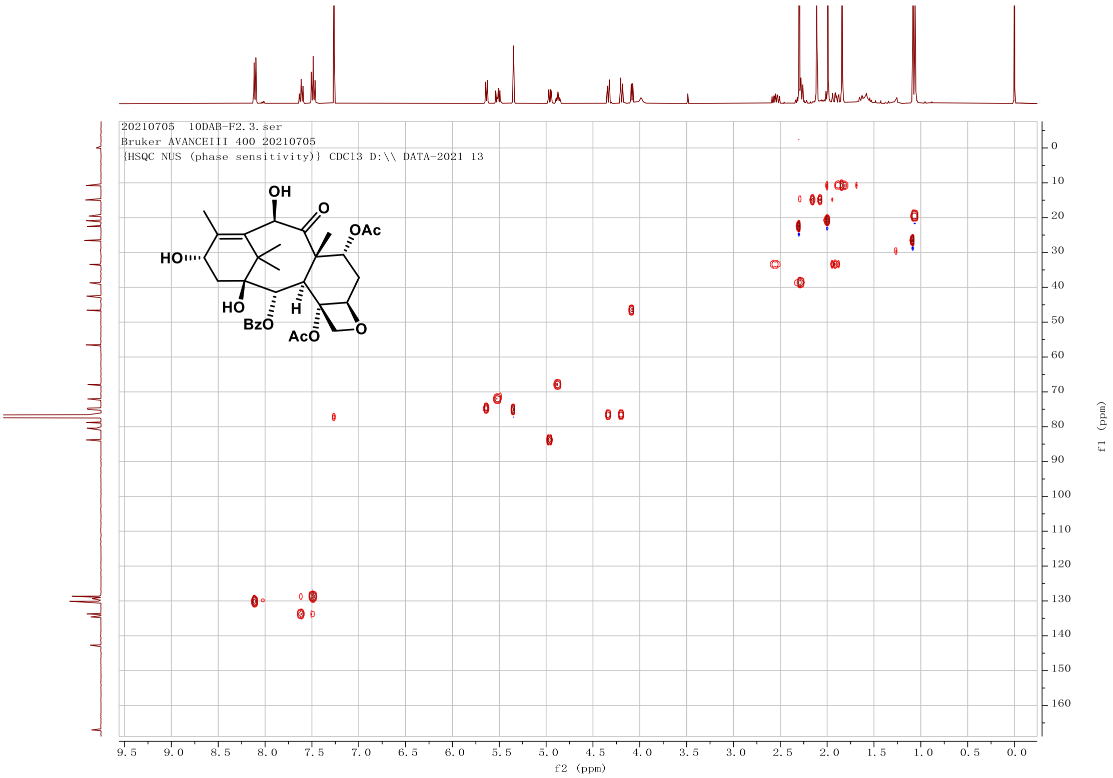

**Fig. S19. HSQC spectrum of 11 in $CDCl_3$.**

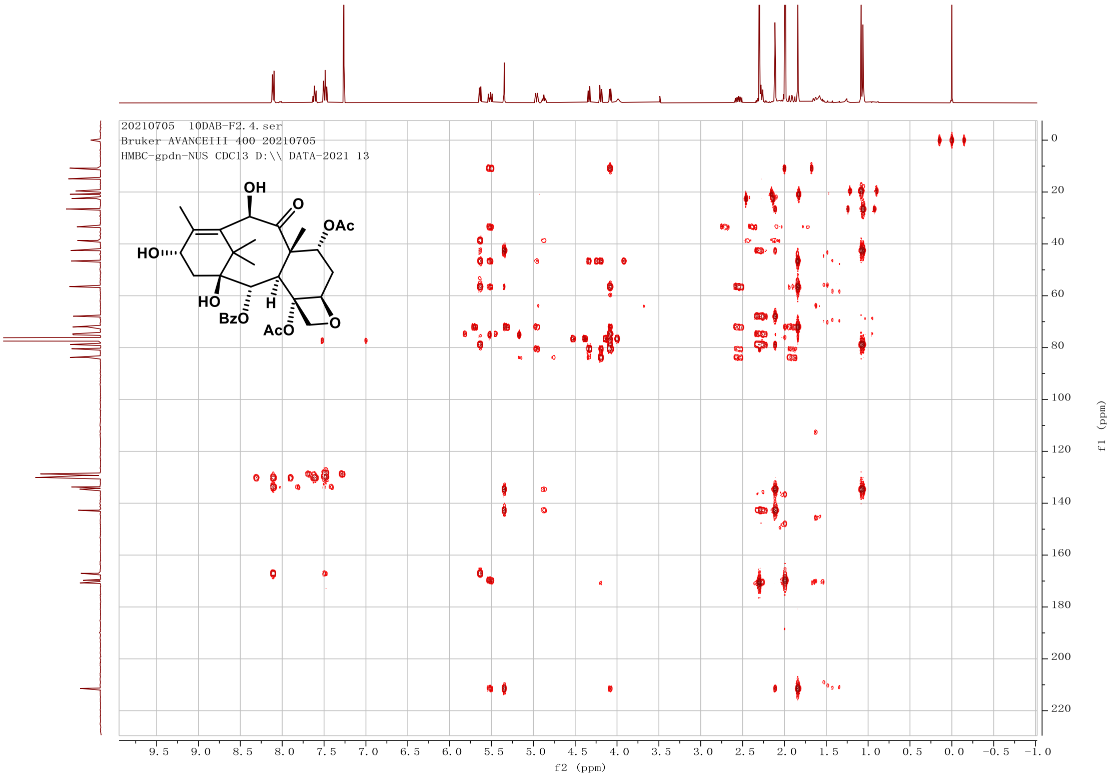

**Fig. S20. HMBC spectrum of 11 in $CDCl_3$.**

Thermo Qexactive Focus Report

compound NO. : F1-ZF
Method : LCMS(compound)-low

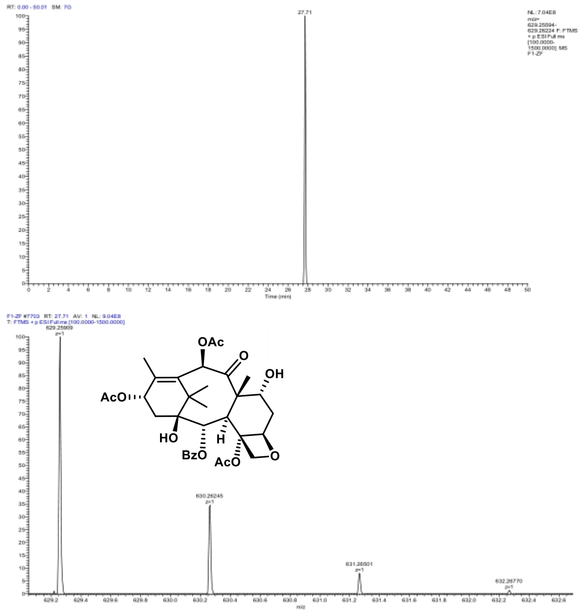

| m/z | Theo. Mass | Delta (ppm) | RDB equiv. | Composition | |
|---|---|---|---|---|---|
| 629.25909 | 629.25925 | -0.26 | 13.5 | C33 H41 O12 | M+H |

**Fig. S21. HR-ESI-MS spectrum of 12.**

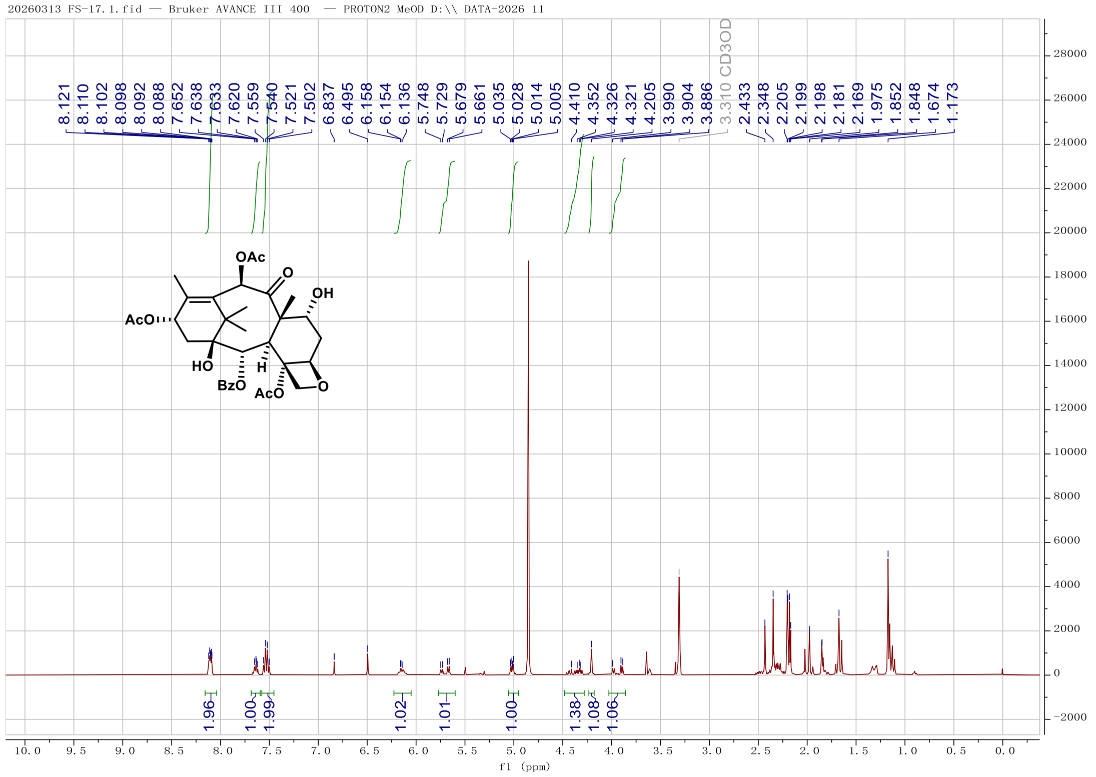

**Fig. S22. [1]H-NMR spectrum of 12 ($CD_3OD$, 400 MHz).**

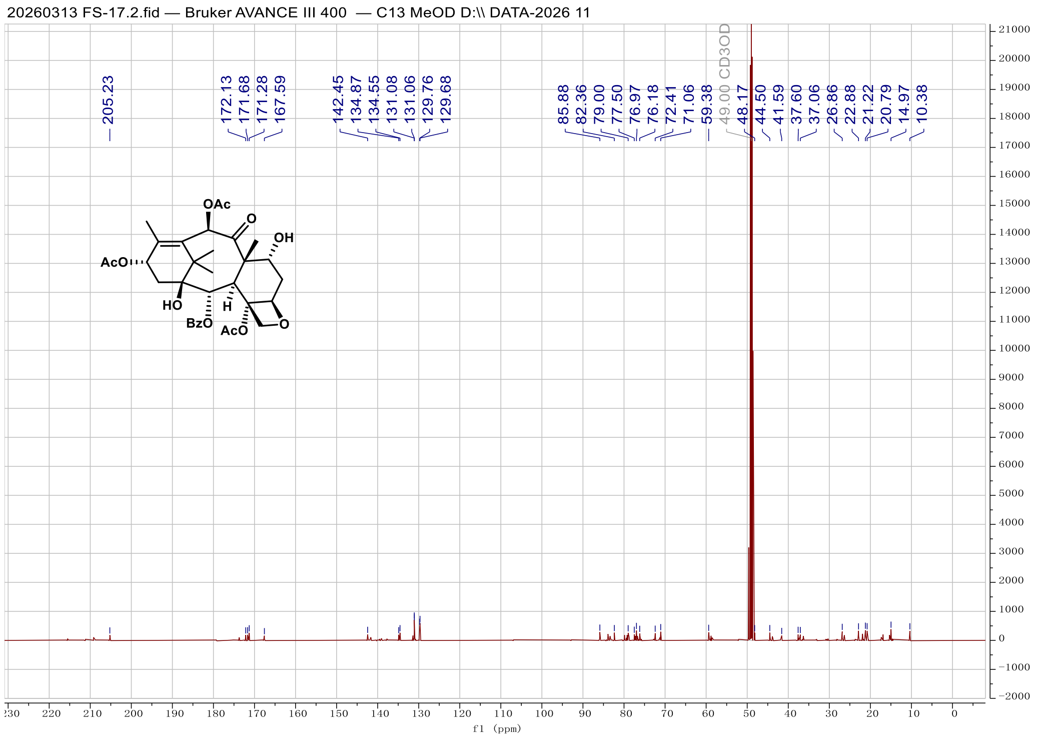

**Fig. S23. [13]C-NMR spectrum of 12 ($CD_3OD$, 100 MHz).**

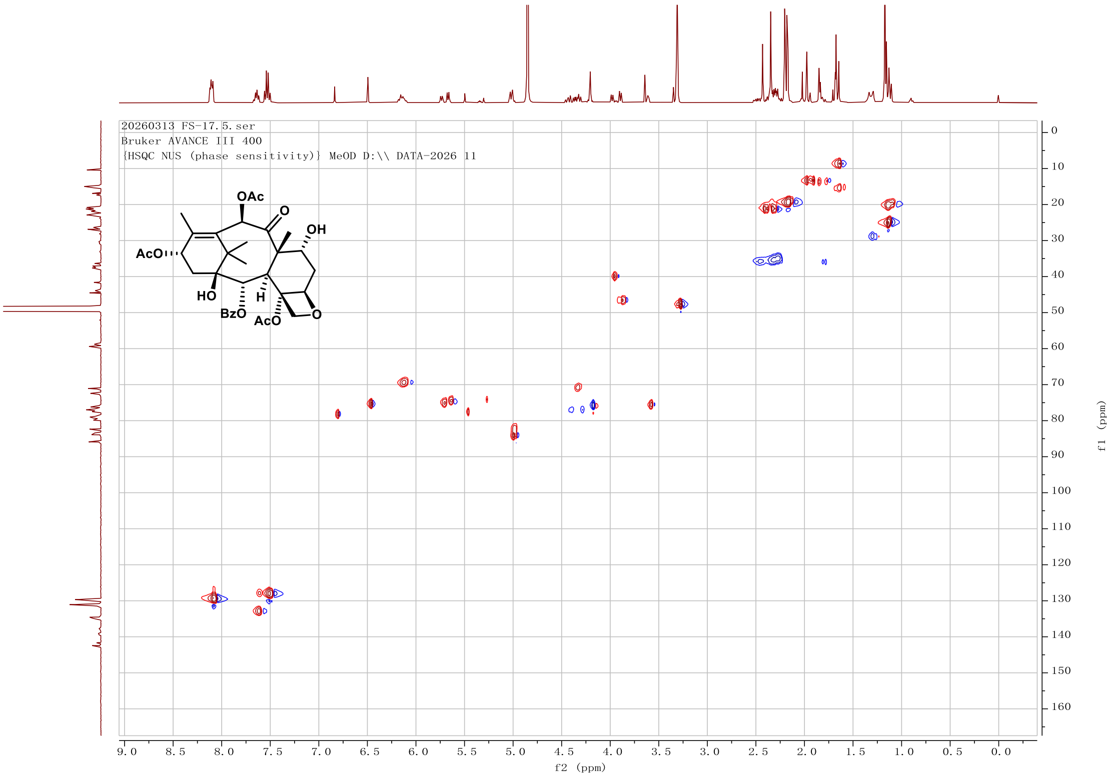

**Fig. S24. HSQC spectrum of 12 in $CD_3OD$.**

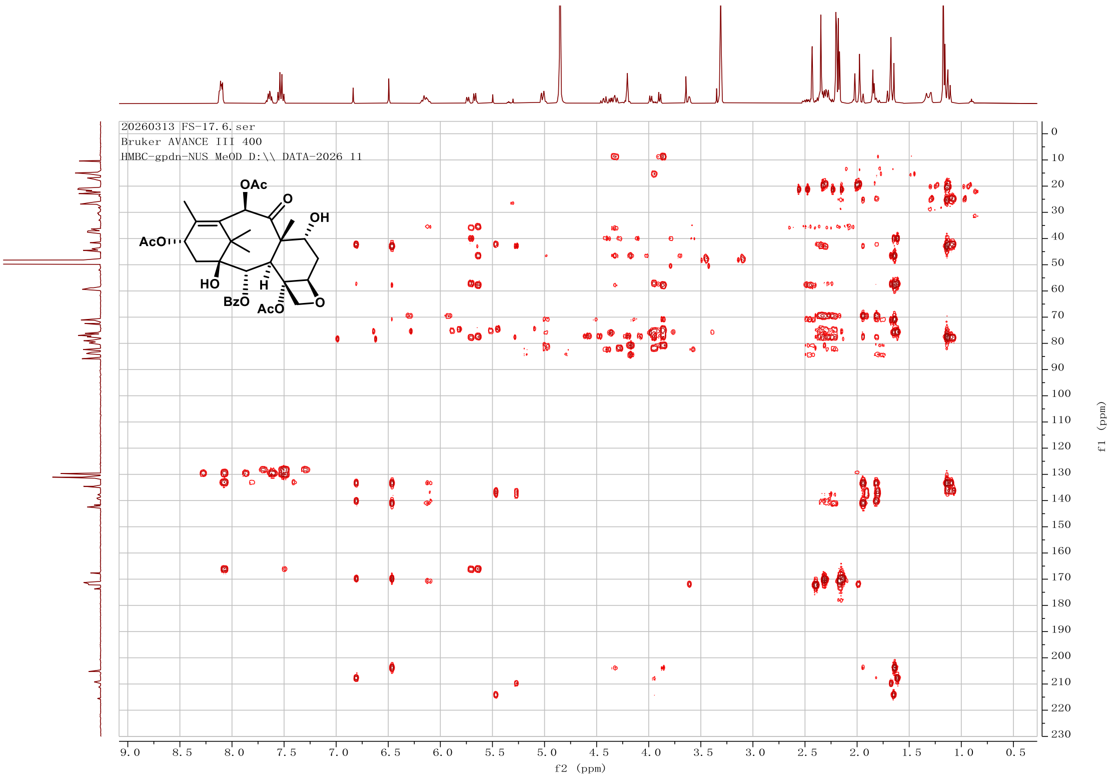

**Fig. S25. HMBC spectrum of 12 in $CD_3OD$.**

Thermo Qexactive Focus Report

compound NO. : FS

Method : LCMS(compound)-low

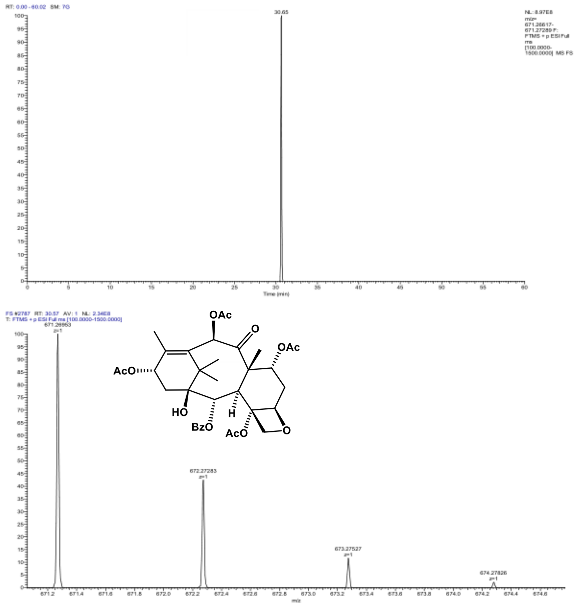

| m/z | Theo. Mass | Delta (ppm) | RDB equiv. | Composition | |
|---|---|---|---|---|---|
| 671.26953 | 671.26982 | -0.43 | 14.5 | C35 H43 O13 | M+H |

**Fig. S26. HR-ESI-MS spectrum of 13.**

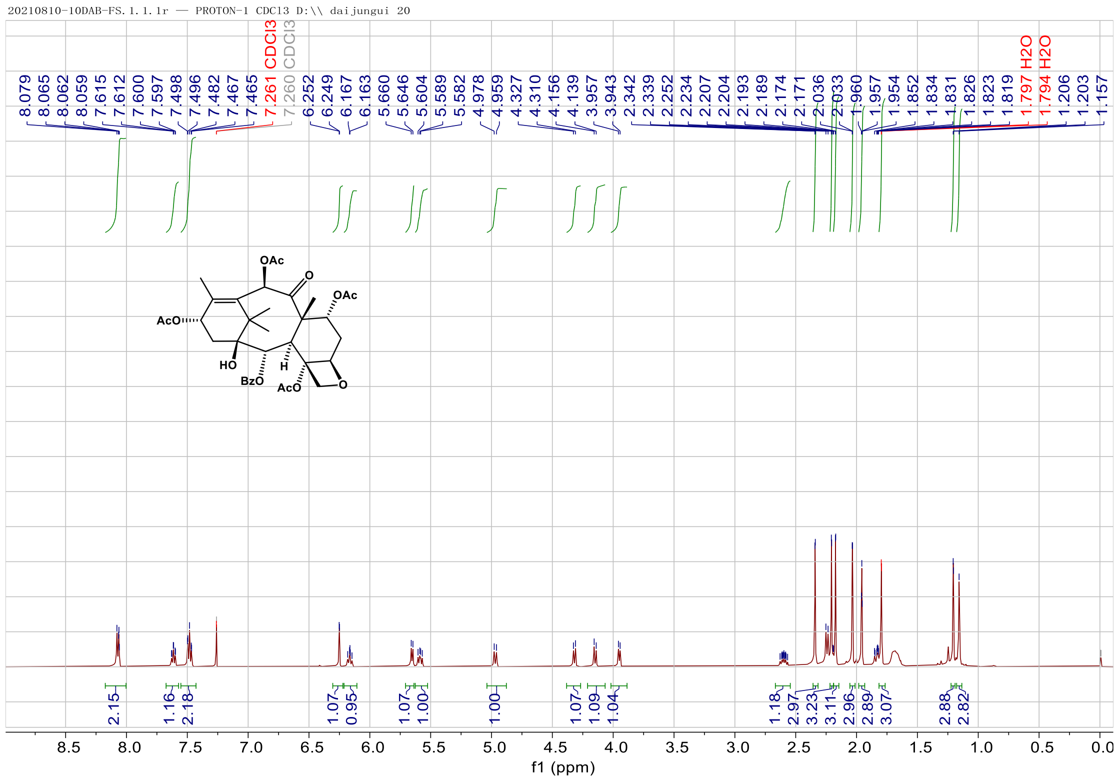

**Fig. S27. $^{1}$H-NMR spectrum of 13 ($CDCl_3$, 400 MHz).**

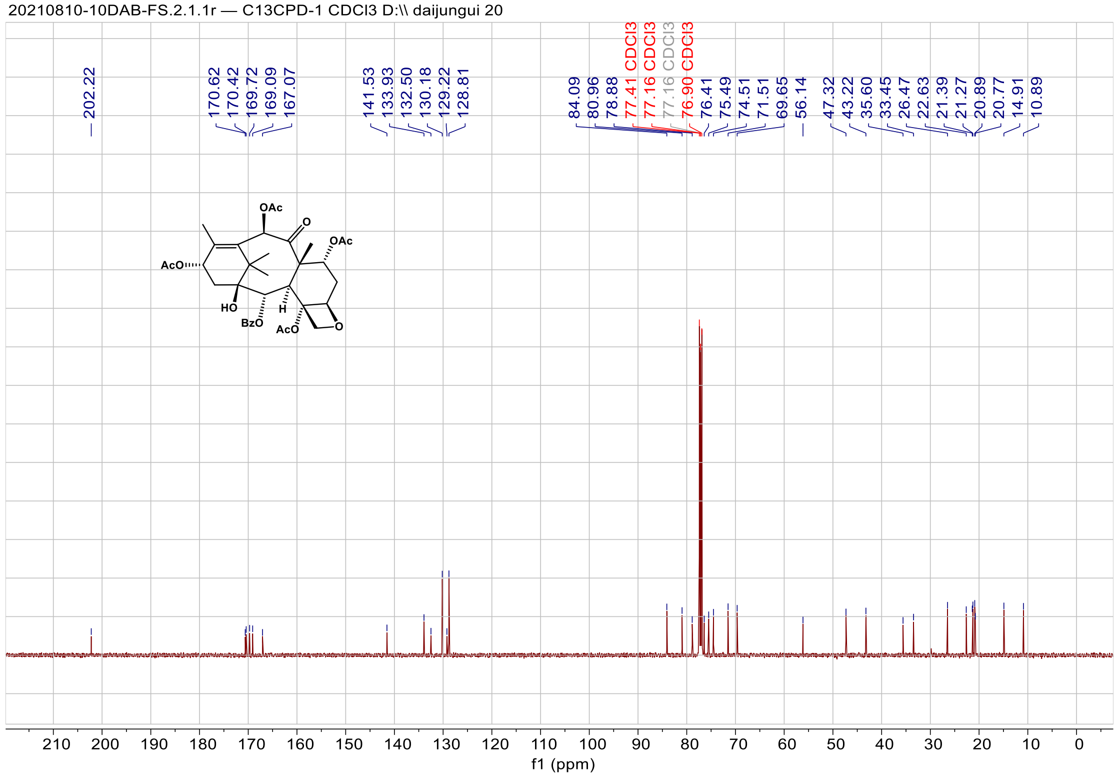

**Fig. S28. $^{13}$C-NMR spectrum of 13 ($CDCl_3$, 100 MHz).**

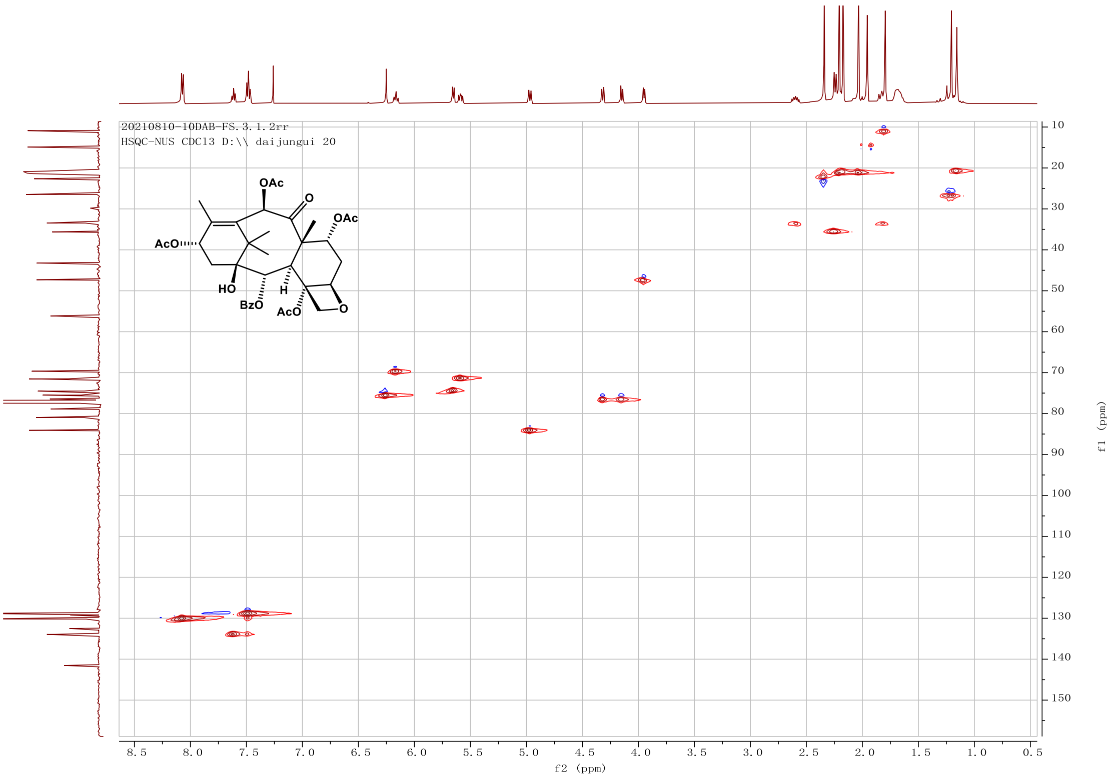

**Fig. S29. HSQC spectrum of 13 in $CDCl_3$.**

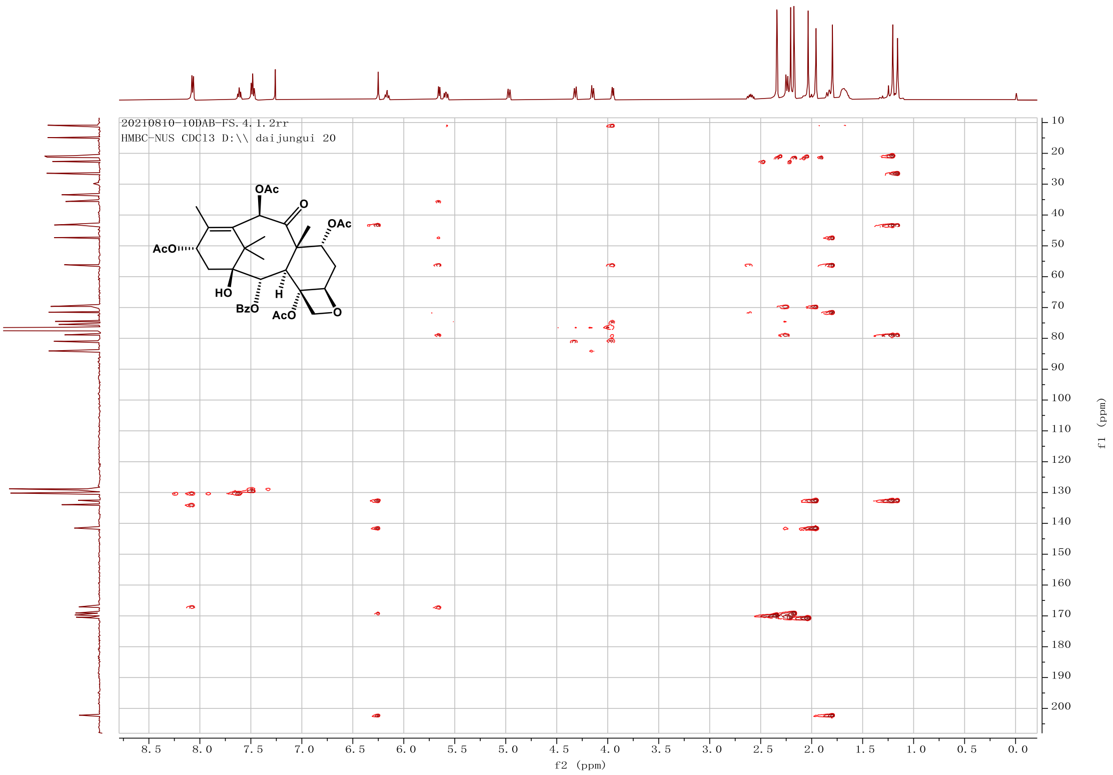

**Fig. S30. HMBC spectrum of 13 in $CDCl_3$.**

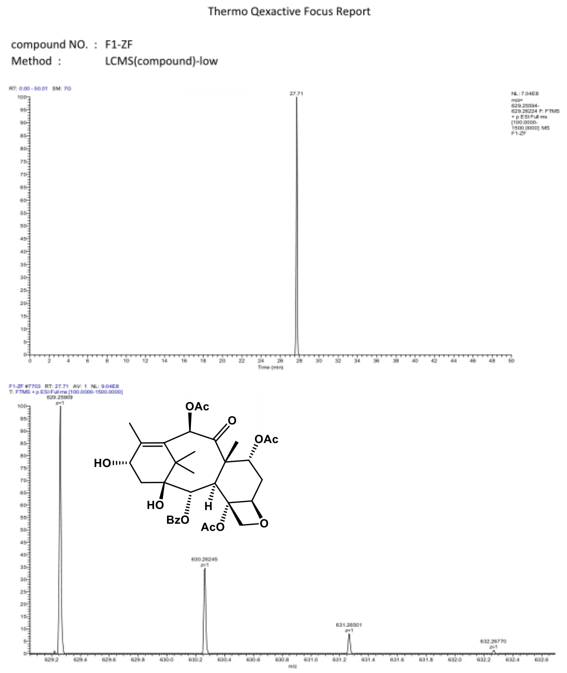

| m/z | Theo. Mass | Delta (ppm) | RDB equiv. | Composition | |
|---|---|---|---|---|---|
| 629.25909 | 629.25925 | -0.26 | 13.5 | C33 H41 O12 | M+H |

**Fig. S31. HR-ESI-MS spectrum of 14.**

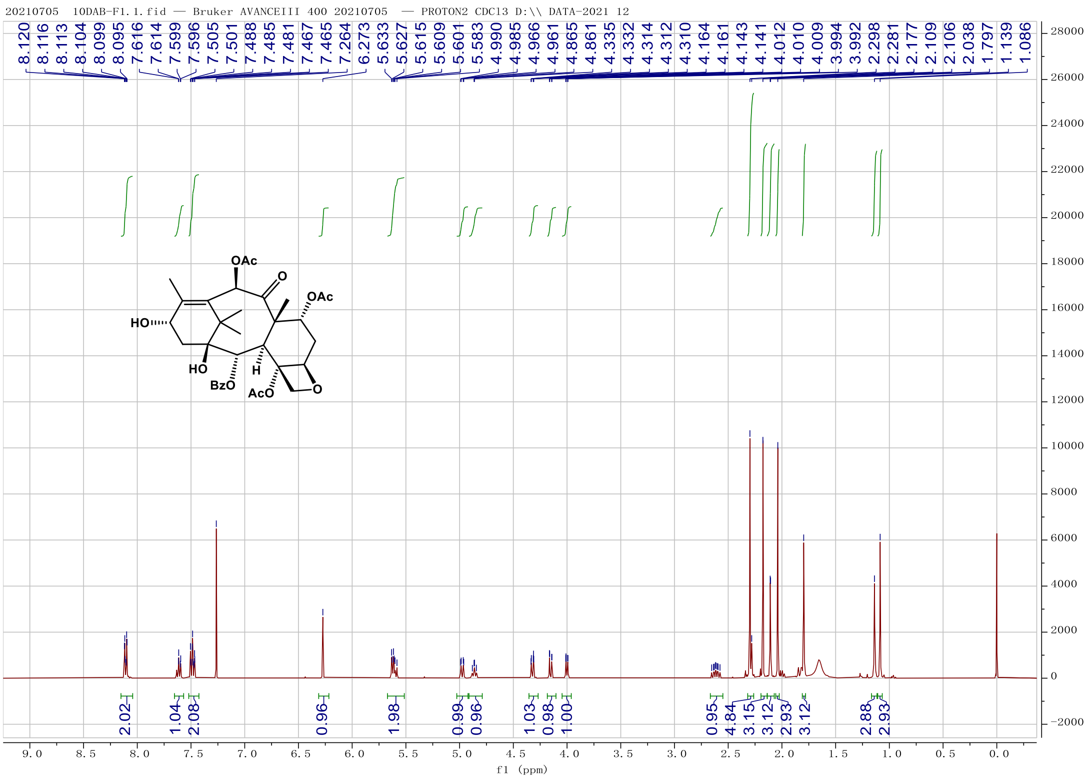

**Fig. S32. $^1$H-NMR spectrum of 14 ($CDCl_3$, 400 MHz).**

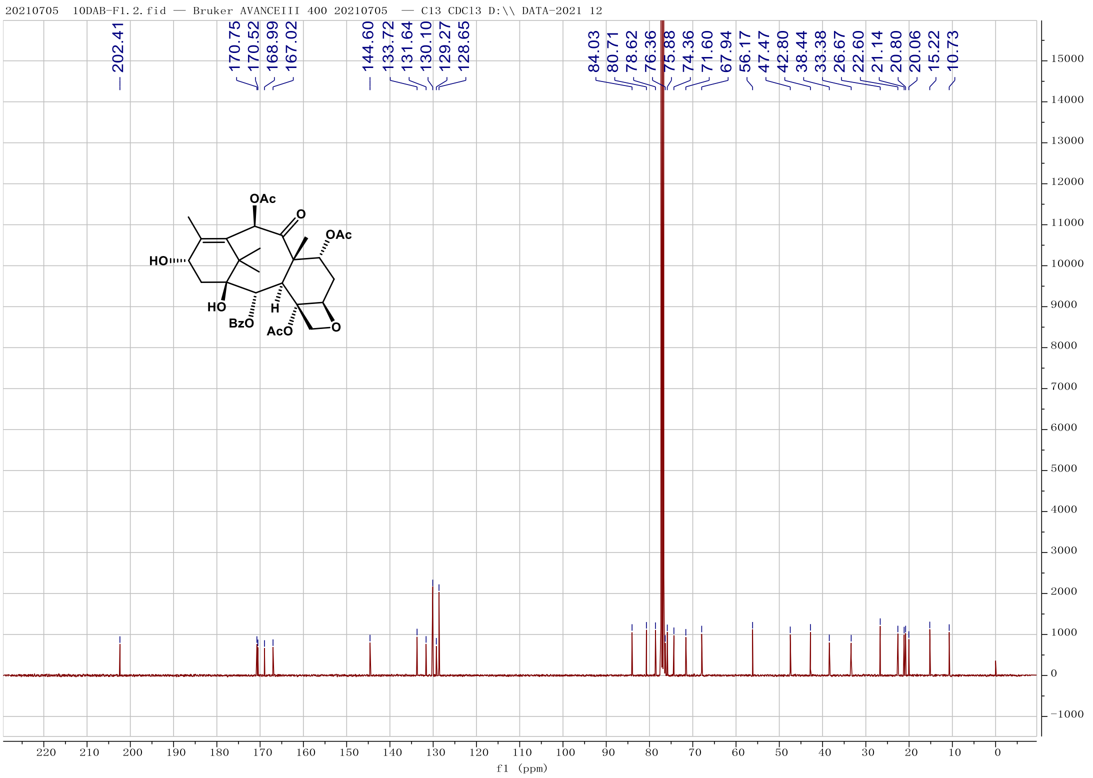

Fig. S33. $^{13}$C-NMR spectrum of 14 ($CDCl_3$, 100 MHz).

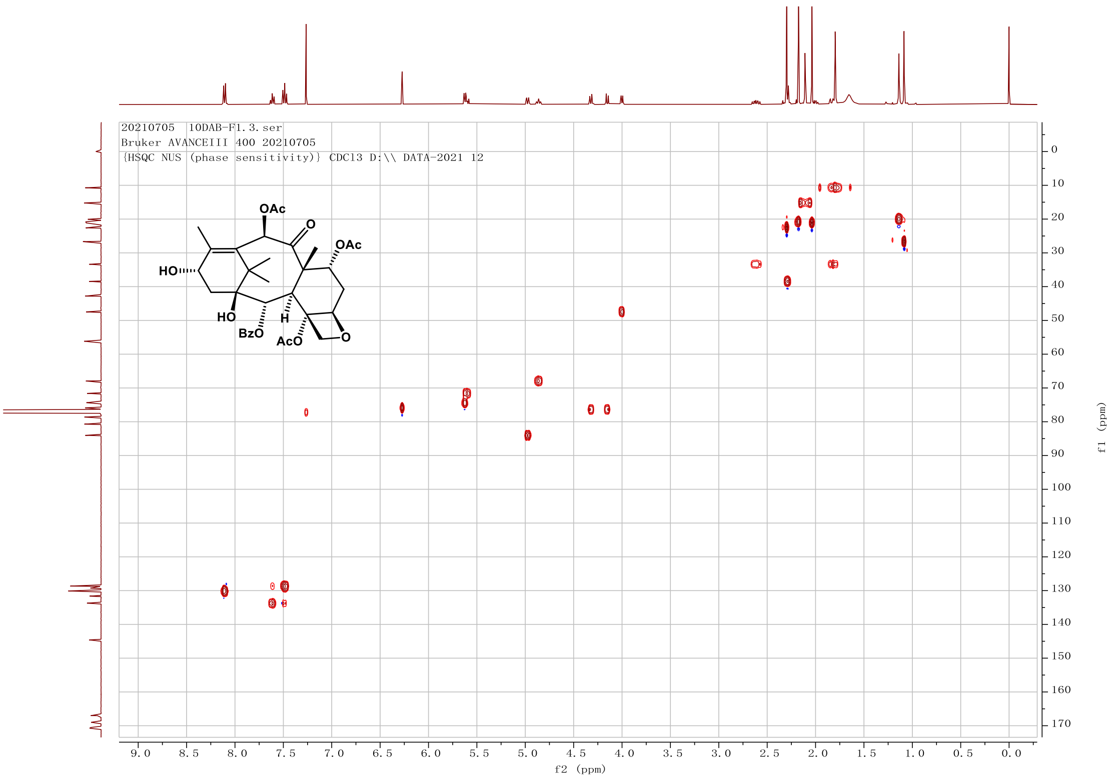

Fig. S34. HSQC spectrum of 14 in $CDCl_3$.

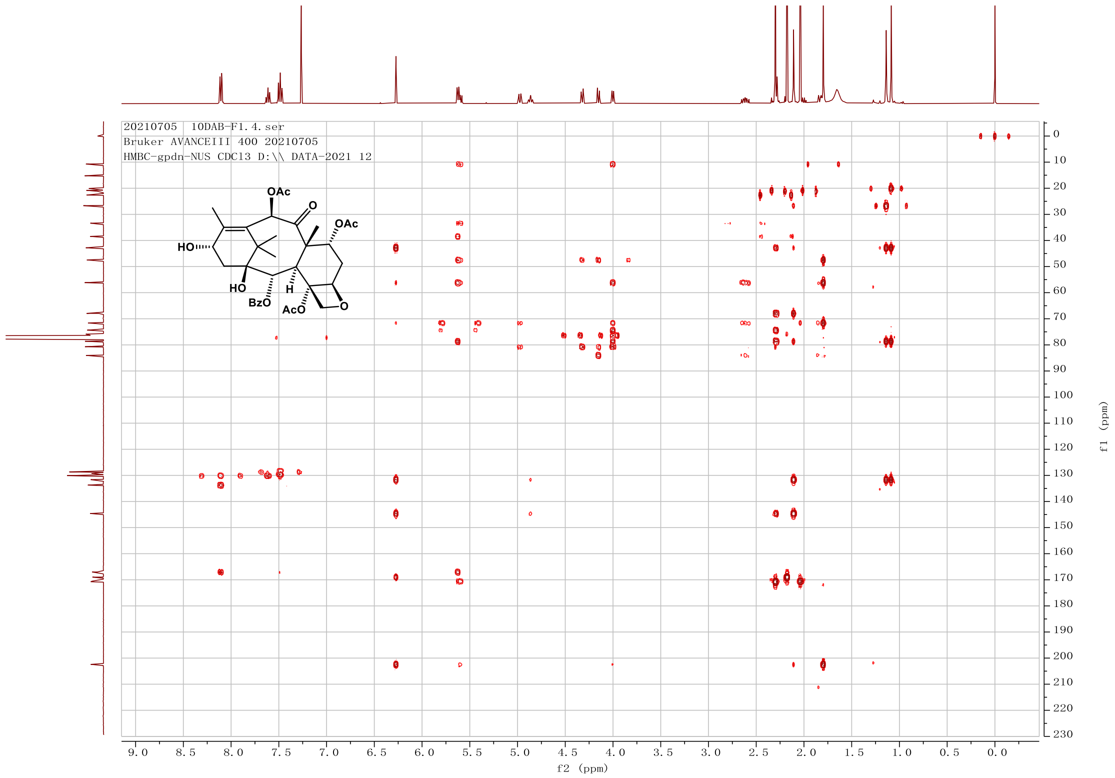

**Fig. S35. HMBC spectrum of 14 in $CDCl_3$.**
